# The impact of dwell time on the contextual effect of visual and passive lead-in movements

**DOI:** 10.1101/2024.10.25.620176

**Authors:** Laura Alvarez-Hidalgo, David W. Franklin, Ian S. Howard

## Abstract

Contextual cues arising from distinct movements are crucial in shaping control strategies for human movement. Here, we examine the impact of visual and passive lead-in movement cues on unimanual motor learning, focusing on the influence of “dwell time”, where two-part movements are separated by the interval between the end of the first movement and the start of the second. We used a robotic manipulandum to implement a point-to-point interference task with switching opposing viscous curl-fields in male and female human participants. Consistent with prior research, in both visual and passive lead-in conditions, participants showed significant adaptation to opposing dynamics with short dwell times. As dwell time increased for both visual and passive signals, past movement information had less contextual influence. However, the efficacy of visual movement cues declined more rapidly as dwell times increased. At dwell times greater than 800 ms, the contextual influence of prior visual movement was small, whereas the effectiveness of passive lead-in movement was found to be significantly greater. This indicates that the effectiveness of sensory movement cues in motor learning is modality dependent. We hypothesize that such differences may arise because proprioceptive signals directly relate to arm movements, whereas visual inputs exhibit longer latency and, in addition, can relate to many aspects of movement in the environment and not just to our own arm movements. Therefore, the motor system may not always find visual movement cues as relevant for predictive control of dynamics.

**News & Noteworthy:** This research uncovers, for the first time, how visual and proprioceptive sensory cues affect motor learning as a function of the pause or “dwell time” in two-part movements. The study has shown that visual lead-in movement cues lose their effectiveness sooner than passive lead-in movement cues as dwell time increases. By revealing the modality-dependent nature of sensory information, this study enhances our understanding of motor control and opens new possibilities for improving therapeutic interventions.

## Introduction

To perform effective arm movements, the human motor system must adapt to both the arm’s dynamics and its environment. Numerous studies have investigated how the motor system deals with changes in dynamics. Viscous curl-fields, in particular, have proven to be useful for examining the adaptation of the human motor system to novel dynamics (1–3). In untrained participants, a curl-field initially disrupts straight hand movements by exerting a force perpendicular to the hand motion and proportional to its velocity. However, adaptation to curl-field dynamics occurs rapidly, usually over a few dozen movement trials. After this period, movement trajectories almost resemble those executed when no curl-field is present. This adaptation, however, is hindered when the direction of the curl-field changes unpredictably. When individuals experience two opposing viscous curl-fields during movements, the learning from the second can disrupt the memory of the first. This phenomenon is known as retrograde interference (4, 5). Similarly, prior learning can affect subsequent learning, a phenomenon known as anterograde interference (6–8). When curl-fields with opposing directions are presented randomly and in balanced proportions, during so-called interference tasks, it has been observed that learning does not occur unless contextual cues are present (9–14).

This interference paradigm provides a valuable technique for testing the impact of contextual information. With effective contextual cues, learning to adapt to opposing curl-fields becomes possible. Prior lead-in movements have been shown to significantly influence the learning of novel dynamics (15–19). Their influence has also been demonstrated in bimanual tasks, extending the understanding of lead-in movements across both arms (20, 21). Notably, in the unimanual case, visual, passive, and active lead-in movements all exert a substantial contextual effect on the adaptation to novel dynamics in subsequent movements (15).

The generalization characteristics of unimanual lead-in movements have been extensively studied across various modalities. These studies include examining the effects of movement direction in visual (18, 22), passive (16), and active lead-ins (17). In terms of prior movement direction, active and passive lead-ins show similar angular generalization characteristics, whereas those of visual lead-ins are broader and weaker. Furthermore, the examination of generalization for additional kinematic movement characteristics, such as distance, speed, or duration, reveals notable differences between visual and passive lead-in movements (19). Another related study found that increased variability in active lead-ins slowed motor adaptation, whereas variations in visual lead-ins had no significant effect (23). It was suggested that this phenomenon could be explained by the differences between their respective generalization characteristics. Taken together, these variations indicate that the way the motor system receives information from past movements significantly affects the formation of motor memory.

It has been shown that the contextual effect of active lead-in movements diminishes as dwell time increases, disappearing almost completely over 1000 milliseconds (15). Given that different lead-in modalities exhibit varying generalization characteristics, this suggests that the influence of dwell time on adaptation might also vary. Although the impact of dwell time in active lead-ins has been studied in some detail, the behaviors of passive and visual lead-ins under similar conditions have not yet been fully examined. In this study, we investigate the impact of dwell time on the contextual effects of visual and passive lead-ins in the learning of opposing dynamics, examining how these effects diminish as dwell time increases. Our study adheres to the same experimental protocol as the previous active lead-in movement study of the effect of dwell time (15), including the use of a vBOT robotic manipulandum (24), in order to facilitate a direct comparison with the results of this prior study.

## Materials and Methods

### Participants

We recruited 48 human participants from the same demographic, each randomly assigned to one of two distinct experiments, with three different dwell time conditions. The first group, consisting of 24 participants (14 females and 10 males, average age: 20.83 ± 1.99 years), participated in Visual Lead-in Experiment 1. The second group, consisting of 24 participants (10 females and 14 males, average age: 21.92 ± 3.23 years), took part in Passive Lead-in Experiment 2. To avoid bias, participants were naïve to the purpose of the experiment. Each condition included eight participants, based on sample sizes used in a previous study (15).

According to the Edinburgh Handedness Questionnaire (25), all participants were right-handed. Prior to their involvement, participants were not informed of the study’s specific objectives, and each participant provided informed written consent. All experiments were conducted in full accordance with a protocol approved by the Science and Engineering Faculty Research Ethics and Integrity Committee at the University of Plymouth, and we strictly adhered to the approved guidelines.

### Apparatus

All experiments were conducted using a vBOT planar robotic manipulandum (24), paired with a 2D virtual reality system, as illustrated in Fig. 1. The handle’s location was determined using optical encoders, sampled at 1000 Hz. Appropriate state-dependent endpoint forces were applied to the handle by controlling the torque of the robot’s drive motors.

**Figure 1.**
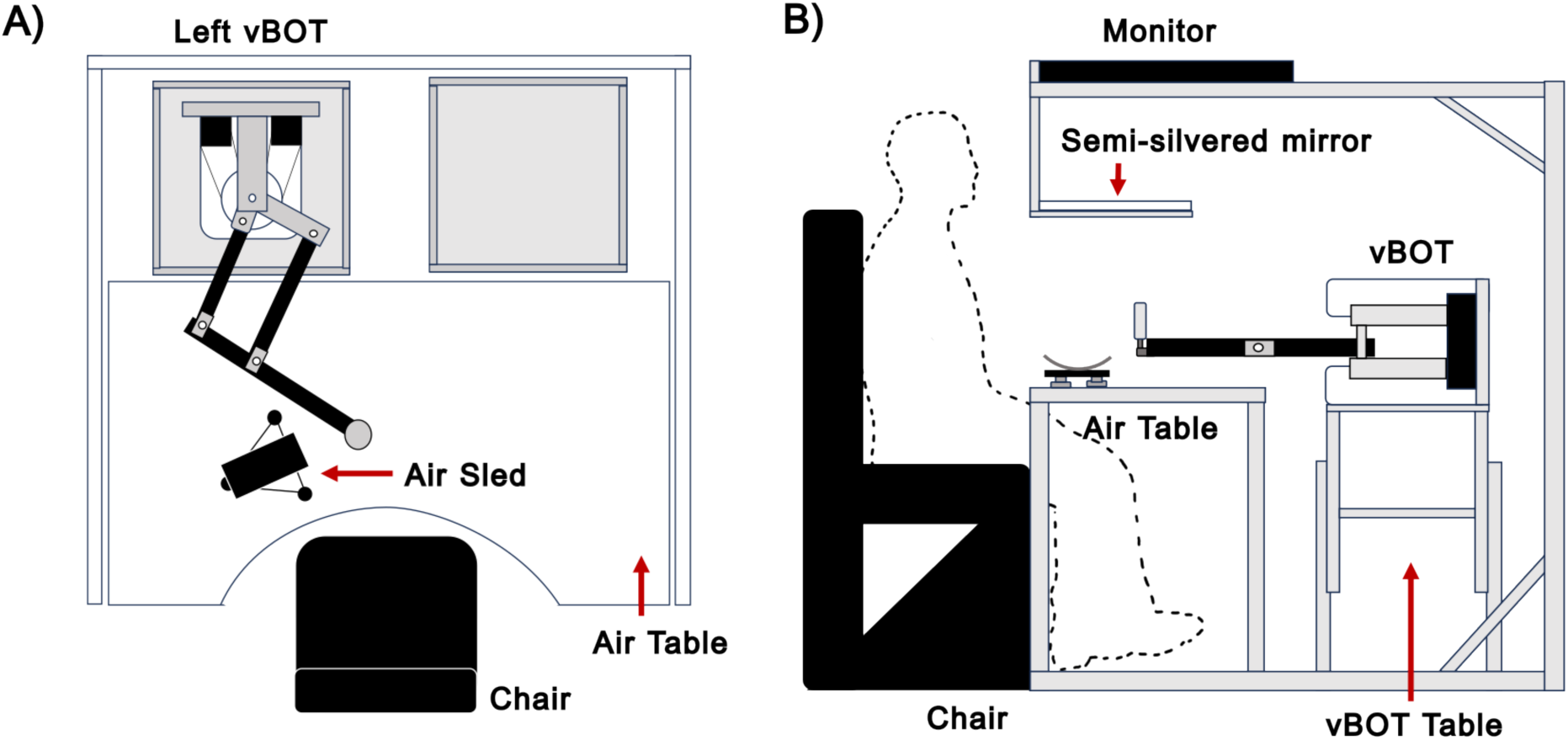
Schematic Illustration of the Experimental Setup. **A:** Plan view of the vBOT robotic manipulandum, air table, and air sled. **B:** Side view of the vBOT robotic manipulandum, showing its virtual reality environment.

A force transducer (Nano 25; ATI), mounted beneath the handle of the robotic manipulandum, measured the applied forces. The output signals were low-pass filtered at 500 Hz using 4^th^-order analog Bessel filters prior to digitization. To minimize body movement, participants were seated in a sturdy chair positioned in front of the apparatus and securely fastened to the backrest with a four-point seatbelt.

During the experiment, participants held the robot handle with their right hand, while their forearm was supported by an air sled, which restricted arm movements to the horizontal plane. Participants could not see their arm or hand directly; instead, they were provided with accurate visual feedback using the virtual reality system. Images of the starting point, via-point, and target locations (each with a 1.25 cm radius), as well as a hand cursor (a 0.5 cm radius red disk), were overlaid to align with the actual plane and location of the participant’s hand. Data were collected at a sampling rate of 1000 Hz and stored on an SSD for offline analysis using MATLAB (The MathWorks Inc., Natick, MA, USA).

### Force Fields

All trials began with a lead-in movement to provide context, followed by an active adaptation movement to a final target. In the adaptation phase, participants made reaching movements in three different types of trials: null field trials, viscous curl force field trials (26, 9), or simulated channel trials (27–29). The velocity-dependent viscous curl-field was implemented according to Equation (1):

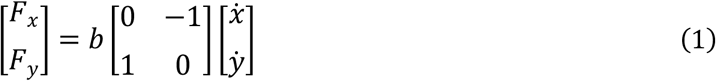

In this equation, the field constant b had a value of ±13Ns/m, where its sign determined the curl-field direction (clockwise or counterclockwise). Each participant was subjected to both directions of the curl-field, which were consistently linked with a specific lead-in movement direction. To minimize potential directional bias, the association between the contextual movement direction and the curl-field direction was counterbalanced across participants.

Mechanical channel trials, from the central location to the final target, were used to assess the level of predictive compensation for the curl-fields (27). These channels were implemented using a spring constant of 6,000N/m and a viscous damping constant of 30Ns/m, with their effects applied perpendicularly to the direction of motion throughout the movement. A channel trial was implemented according to Equation (2):

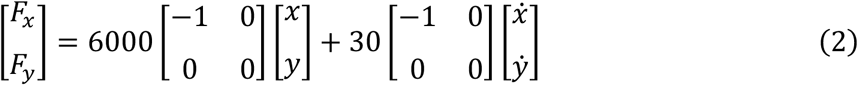

Two separate experiments were conducted with different groups of participants, each using a similar protocol. In each experiment, participants waited for either a visual cue or a passive cue, depending on the experimental setup.

In the Visual Lead-in Experiment 1, the participant’s hand was initially located at the central via-point. Participants observed a cursor moving from a starting position to a via-point while their right hand remained stationary. In contrast, in the Passive Lead-in Experiment 2, the participant’s hand was initially positioned at a peripheral starting point and was pulled to the central via-point by the robotic system, thus avoiding active movement generation.

Following the cue, participants were instructed to continue performing an active movement only after a predefined dwell time had elapsed. This target dwell time was nominally set at either 150 ms, 300 ms, or 500 ms, depending on the experimental condition. These constituted the minimum dwell times that could be accepted in each respective condition. In both experiments, the active movement consisted of a 12 cm motion to the target.

### Minimum Jerk Trajectory Generation

The contextual lead-in motions followed a trajectory generated by the vBOT system, rather than by the participant, from the given starting location to the central via-point location. For a one-dimensional movement starting at location x_0_, with duration T and distance D, the minimum jerk position trajectory x(t) at time t, is given by Equation (3).

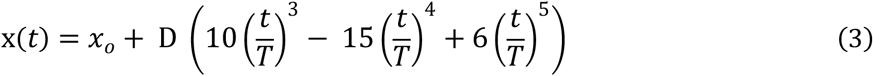

See also (30). The corresponding velocity trajectory v(t) at time t is given by the time derivative of position:

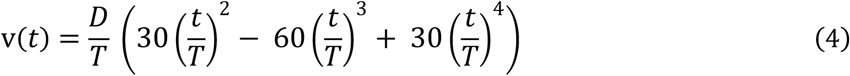

In our experiments, trajectory parameters were specified as T = 663 ms and D = 10 cm, as used in the previous active lead-in study (15), which led to smooth and achievable passive lead-in movements. The termination condition for the lead-in movement also followed those used in the previous study, based on both position and speed, as described in the relevant sections below.

### Visual Lead-in Experiment 1

#### Protocol

A schematic depiction of the experimental protocol experienced by a participant is illustrated in Fig. 2. This outlines the training and testing procedures, highlighting the directions used for both contextual and adaptation movements. In total, there were four different starting points for the contextual movement. These were located on the circumference of a circle with a 10 cm radius, with its center corresponding to the central workspace coordinate position (0, 0). Specifically, the start locations were positioned at angles of 45°, 135°, 225°, and 315°. The endpoint for the contextual movement was always at the via-point, also located at the central workspace coordinate position.

**Figure 2.**
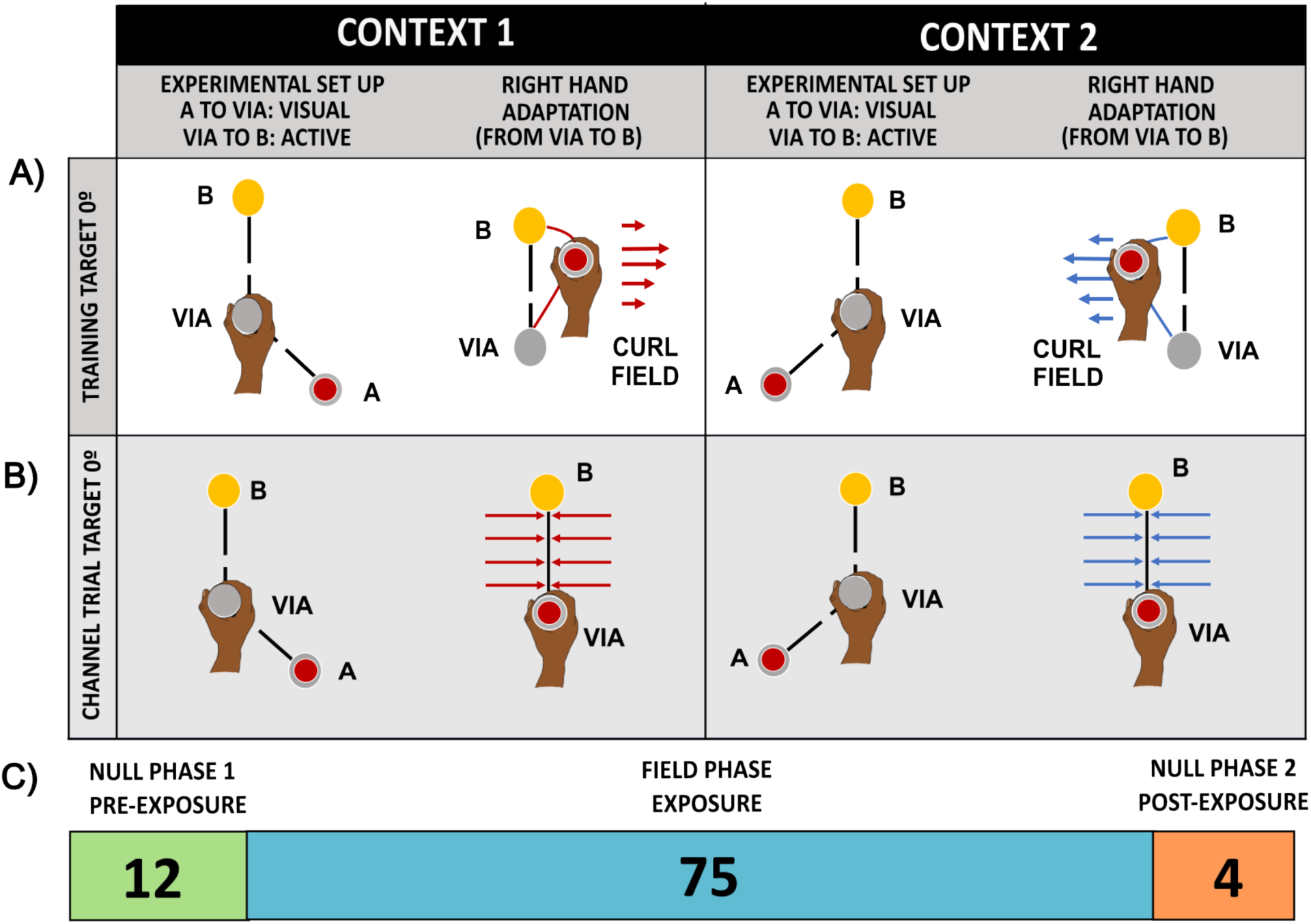
Visual Lead-in Movements Experimental Design. **A**: Null and Training Trials. Here, we show one of the four movement target locations (0°) with both of its visual contextual lead-in conditions, indicating the start, via, and target locations, as well as the hand position and red cursor starting locations. Each trial session began with the participant’s hand at the via-point. A visual lead-in movement then followed from the peripheral start position to the via-point, and after the appropriate dwell time, the participant moved to the displayed target position. **B:** Channel Trials. These trials only used a single adaptation movement direction to a target point at 0°, which was associated with two visual lead-in directions, one for each curl-field direction, with starting points at 135° and 225°. **C.** Outline of the trials in the visual lead-in experimental protocol.

For the adaptation movement, there were four target points, located on the circumference of a circle with a 12 cm radius with its center again corresponding to the central workspace coordinate position. This time, the target locations were positioned at angles of 0°, 90°, 180°, and 270°. Each adaptation movement direction was associated with two distinct contextual movement directions. These relationships determined the direction of the viscous curl-field experienced by participants during the adaptation phase of a trial. This resulted in eight unique combinations of contextual and adaptation movements, as outlined in Table 1. Channel trials were conducted exclusively for movements toward the 0° target. The corresponding lead-in contextual movements originated from locations at either 135° or 225°.

**Table 1.** Specification of contextual and adaptation movements and curl-field directions. The table provides the specific combinations of adaptation target angles and contextual starting point angles used in the experiments. It also shows their corresponding relationship between these angles and the direction of the curl-field. In a given experiment and dwell time condition, the label ‘Normal’ indicates the curl-field direction for four participants, and ‘Reverse’ represents the direction for the remaining four participants. This counterbalancing approach was used to minimize potential biases that might arise from the interaction between the direction of movement and the curl-field.

| Adaptation target | Contextual start | Field direction<br>Normal | Field direction<br>Reverse |
| --- | --- | --- | --- |
| 0° | 135° | CCW | CW |
| 0° | 225° | CW | CCW |
| 90° | 315° | CCW | CW |
| 90° | 225° | CW | CCW |
| 180° | 45° | CCW | CW |
| 180° | 315° | CW | CCW |
| 270° | 135° | CCW | CW |
| 270° | 45° | CW | CCW |

Each trial began with the participant holding the robotic handle and pressing the activation switch. They were then pulled by the robotic arm to the handle’s starting location for the trial, which in this case was the via-position. At this stage, a grey disk was shown at the starting point of the contextual movement. Once the handle remained stationary within its via-point location for 300 ms, the trial proceeded. Then, grey and yellow disks, representing the via-point and target point, respectively, appeared simultaneously with a red moving cursor, which provided the contextual visual movement.

The red cursor moved in a minimum-jerk trajectory directly along the lead-in path towards the via-point. The lead-in terminated when the cursor reached within a predefined tolerance of the via-point goal location. In the visual experiment, since the lead-in cursor movement was unaffected by the participant’s behavior, a tight tolerance was used, corresponding to the cursor moving within 0.5 cm of the central via location. This resulted in a theoretical lead-in duration of 513 ms. The vBOT recorded a flag to indicate when the visual lead-in movement had started and terminated to facilitate later offline MATLAB analysis of the data.

When the contextual movement was complete, its starting location changed from grey to white, accompanied by an auditory beep. After they perceived that the predefined dwell time had been met, participants initiated an active movement to the target. If they attempted to move prematurely or failed to commence the movement within the required duration, the vBOT aborted the trial. This ensured that after the contextual movement, participants remained at the via-point for the necessary dwell time before making the adaptation movement. The trial terminated when the participant actively moved the cursor into the yellow target and stayed there for 300 ms.

### Experimental Block Design

An experiment consisted of blocks of 18 trials, each comprising 16 field trials and 2 clamp trials. During the field trials, all the four targets were presented twice, in both opposing field directions. Each experiment began with 12 blocks of null field trials at the specified experimental dwell time relationship, allowing participants to acclimatize to making movements using the robotic manipulandum. This was followed by a training session consisting of 75 blocks of velocity-dependent curl-fields, in which the field direction was determined by the direction of the contextual movement. The experiment concluded with 4 blocks of washout trials, to examine the aftereffects of the curl-fields being removed. Channel trials, set at the training timing relationship, were interspersed throughout the experiment.

In total, there were 1,638 trials. The study lasted around three hours, although some participants took longer in experimental conditions with longer dwell times. Participants were required to take short rest breaks every 195 to 205 trials. The exact number was determined randomly so it wouldn’t lead to a trend in the data across participants at break boundaries. They could also rest at any time. The timeline of the trials in the experiment is shown in Fig. 2C.

To examine the effect of different dwell times on curl-field adaptation, groups of eight different participants performed the experiment in three different goal dwell time conditions: short (150 ms), medium (300 ms), or long (500 ms). These were selected based on an interpretation of the results from the previous active dwell time study (15), given the hypothesis that the decline in compensation for passive and visual lead-ins would likely follow a similar trend. These earlier results suggested that there would be a substantial reduction in the compensation effect with a requested dwell time of 500 ms. We also decided it would be useful to include additional data points for shorter dwell times, so we selected 150 ms and 300 ms. We did not include a very short dwell time condition, since we expected this would simply yield a high level of compensation across both the visual and passive experiments, and that it would be more informative to sample dwell times at points where greater reductions in compensation were anticipated. Finally, we note that dwell times longer than 500 ms substantially increase the duration of the experiments, making them more challenging to run.

The actual dwell times performed by participants were calculated from the recorded movement data trajectories, and these values were used in subsequent data analysis.

### Passive Lead-in Experiment 2

#### Protocol

The Passive Lead-in Experiment 2 was like the Visual Lead-in Experiment 1 with the exception that the lead-in movement modality is passive movement rather than visual, requiring a corresponding change in the hand’s starting position for each trial. A schematic depiction of the passive lead-in experimental protocol is illustrated in Fig. 3 for a single target direction.

**Figure 3.**
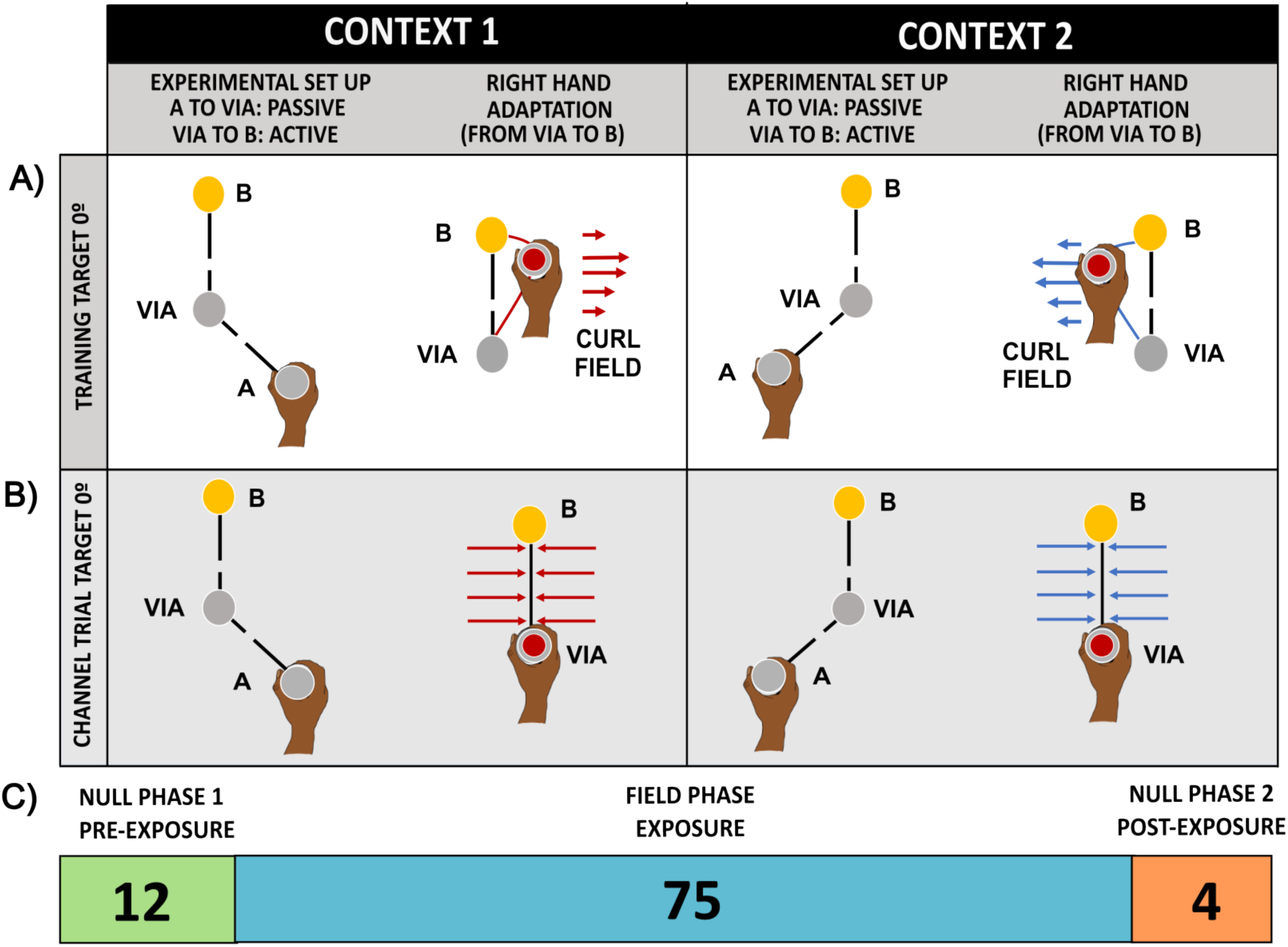
Passive Lead-in Movements Experimental Design. **A**: Null and Training Trials. Here, we show one of the four movement target locations (0°) with both of its passive contextual lead-in conditions, indicating the start, via, and target locations, as well as the hand position and red cursor starting locations. Each trial session began with the participant’s hand at the start point. A passive lead-in movement then followed from the peripheral start position to the via-point, and after the appropriate dwell time, the participant moved to the displayed target position. **B:** Channel Trials. These trials only used a single adaptation movement direction to a target point at 0°, which was associated with two visual lead-in directions, one for each curl-field direction, with starting points at 135° and 225°. **C.** Outline of the trials in the passive lead-in experimental protocol.

To initiate a trial in the passive experiment, the vBOT moved the participant’s hand to the starting location of the lead-in movement. At this stage, a grey disk was shown only at the starting point of the contextual movement. Once the handle remained stationary within this starting location for 300 ms, the trial proceeded. Then, grey and yellow disks appeared to identify the via-point and target point, respectively.

The robot’s handle then physically moved the participant’s hand towards the via-point, following a minimum jerk trajectory, via a stiff spring (k = 2,000 N/m), thereby generating a passive lead-in movement. No visual cursor movement was shown during this operation to avoid the generation of visual cues.

The passive lead-in was successfully completed when the handle reached a position within a predefined tolerance of the via-point goal location. Here, as the lead-in movement could be affected by the participant’s behavior, a more relaxed termination tolerance was needed, corresponding to moving within a distance of 1.5 cm from the central via location while maintaining a speed below 5 cm/s. Provided that the participant did not hinder handle movements, this resulted in an unperturbed theoretical lead-in duration of 559 ms; however, the actual value varied slightly if they resisted or assisted movement.

Once the contextual movement was complete, the via-point turned white, and a red cursor appeared, indicating the participant’s hand position. Upon perceiving that the required dwell time had elapsed, participants initiated their movement towards the target point. If they began the movement prematurely or failed to initiate the active movement within the stipulated duration, the vBOT intervened, forcing the participant to restart the trial. The trial terminated when the actively moved cursor reached the yellow target disk and stayed there for 300 ms.

## Data Analysis

The experimental data were processed offline using MATLAB R2024a. Learning was evaluated using two metrics: kinematic error during adaptation movements and force compensation during channel trials. Additionally, the actual dwell times were calculated for all participants in each experiment.

### Hand Trajectory Plots

Hand trajectory plots provide a useful indication of participants’ movement behavior, showing the effect of the introduction and subsequent removal of the curl-fields. In both lead-in experiments, there were four critical stages: The last block of null-field exposure, the initial block of curl-field exposure, the final block of curl-field exposure, and the initial washout block after curl-field removal.

At critical stages in the experiment, we selected hand trajectories only for the adaptation phase of each movement, excluding the channel trials. We then analysed each of these four selected blocks of trials separately. In each chosen block, there were eight different combinations of target direction and curl-field direction per participant. We calculated the mean and standard error (SE) across participants.

### Kinematic Error

For each null and viscous curl-field trial, the kinematic error during the adaptation phase of the movement was quantified using its Maximum Perpendicular Error (MPE). The MPE was calculated as the maximum deviation of the hand’s path from the straight line connecting the actual start position of the movement to the center of the target.

For each participant, an average MPE was first computed for all movements for each curl-field direction separately, across two blocks (16 trials for each direction). Then, the mean and standard error (SE) of these signed MPEs were calculated across all participants and plotted to indicate behavior in each field direction.

Next, we appropriately combined the individual participants’ results from clockwise (CW) and counterclockwise (CCW) field trials to obtain the overall mean and standard error (SE) of the MPE values across all participants and for both field directions, giving values corresponding to 32 field trials. In the latter calculation, the sign of the MPE was flipped appropriately so that both force field directions would produce positive errors.

### Estimation of Adaptation to the Curl Field

When making point-to-point movements in a null field condition (i.e., during free unloaded hand movements), movements are typically characterized by straight trajectories. On initial exposure to a curl field, participants are initially pushed off these straight paths by a velocity-dependent perpendicular force, which results in looped paths. However, participants rapidly adapt and generate predictive compensatory forces to counteract the effect of the curl field (9). Predictive feedforward adaptation is a process by which the motor system learns to compensate for changes in the motor task, such as arm dynamics, by making compensatory adjustments independently of feedback (31, 32).

Channel trials, in which movements are constrained to straight paths, provide a means to directly estimate this compensation, and thereby provide a means to estimate the extent participants compensate for the effect of the curl field (33). During channel trials, the perpendicular force exerted on the wall of the simulated channel was measured using a force transducer. This enabled the estimation of predictive feedforward adaptation to the applied curl-field, which was achieved by regressing measured force against the movement velocity along the channel without an offset term (34). The use of a channel is often considered a more reliable method to assess learning than simply observing the reduction of kinematic error during force field adaptation, which can be influenced by muscle co-contraction (35, 36, 33).

To indicate the progressive increase in compensation over the course of the experiment for each of the curl field directions separately, an average curl-field compensation value was first computed for each participant across all movements in each curl-field direction, using data from two blocks (each containing two channel trials per direction). Then, the mean and standard error (SE) of these signed compensation values were calculated across all participants and plotted to indicate behavior in each field direction.

Furthermore, to indicate the overall increase in compensation over the course of the experiment, the curl-field compensation values were averaged over the four consecutive channel trials across two blocks, which corresponded to both field directions. This was achieved by appropriately adjusting the sign of the values before calculating the mean. To examine the overall adaptation to the two viscous curl-fields during training, we then computed the mean and standard error (SE) of compensation for the training trials across all participants.

To estimate the final levels of compensation achieved across the different experimental and dwell time conditions, final compensation values were calculated by averaging sign-adjusted compensation values for both field directions over the last 20 consecutive channel trials (the last 10 exposure blocks) in the exposure phase of the experiment. Finally, the mean and standard error (SE) of compensation for the training trials across all participants were calculated.

### Estimation of Dwell Time in Experiments 1 and 2

To estimate the actual dwell times of movements performed by participants, we calculated the time difference between the end of the lead-in movement and the start of the adaptation movement. The termination of the lead-in movement was identified using a recorded lead-in flag signal supplied by the vBOT data. After the contextual lead-in had terminated, the hand was positioned at the via-point in both the visual and passive lead-in experimental conditions.

The start of the adaptation movement was estimated to occur when the hand position moved away from the central via-point by more than its radius (1.25 cm), placing it just outside the via circle, and was traveling at a speed greater than 5 cm/s. To enhance the robustness of the detection process, the start of the adaptation movement also had to satisfy a minimum dwell time criterion set by the specific dwell time condition. The dwell time formula is shown in Equation (5):

The lead-in duration for each trial was also directly determined from the recorded lead-in flag signal.

### Reanalysis of Previous Active Lead-in Data

In order to compare the results of the current study with relevant results from a previous study (15), which examined the effect of dwell time on active lead-in, we reanalyzed the former dataset using our current MATLAB code. In addition to examining the former active lead-in datasets, we also reanalyzed short-latency passive and visual experimental data from the former study.

The only difference between the analysis of the active lead-in data and the current visual and passive datasets was how the start and end of the lead-in movement were determined. As there was no flag signal indicating the operation of the active lead-in, both the start and termination of the lead-in movement were determined based on the hand position and speed. The start of the lead-in was determined when the hand exceeded a threshold distance (1.25 cm) from the start location and its speed was greater than 5 cm/s. Correspondingly, lead-in was deemed to have terminated when the hand was within a threshold distance (1.25 cm) from the via location and its speed was less than 5 cm/s.

We note that the reanalysis of the short-latency passive and visual experimental data from the former study made use of a lead-in flag signal and was identical to the analysis of the current passive and visual lead-in data.

### Statistical Analysis

Statistical analyses were conducted using JASP (37). To ensure the validity of our analyses, Mauchly’s test was used to assess the assumption of sphericity when applicable. In cases where this assumption was violated, the degrees of freedom were adjusted using the Greenhouse-Geisser correction. The significance level (α) was set at 0.05 for all tests, and omega squared (ω²) was reported as the effect size. Omega squared (ω²) serves as a measure of how much variance in the dependent variable is explained by the explanatory variables.

Each of the two experiments was conducted with different participants under three dwell time conditions, resulting in a total of six sets of MPE data and six sets of force compensation data. For each of these six datasets, we first performed repeated measures ANOVAs to examine differences in MPE and force compensation at various critical phases within each experimental condition.

To evaluate kinematic error, we compared the final pre-exposure MPE (the mean of the last two pre-exposure blocks), the initial exposure MPE (the mean of the first two exposure blocks), the final exposure MPE (the mean of the last two exposure blocks), and the initial washout MPE (the mean of the first two washout blocks). This comparison was conducted using a repeated measures ANOVA, with phase serving as a repeated factor (4 levels). When a significant main effect was observed, we proceeded with paost-hoc comparisons using the Holm-Bonferroni correction.

For force compensation, we compared the final pre-exposure (the mean of the last pre-exposure blocks), the final exposure (the mean of the last two exposure blocks), and the initial washout using a repeated measures ANOVA, with phase (3 levels: pre-exposure, final exposure, and washout) as the repeated factor.

To compare MPE across dwell time conditions within each experiment, we performed ANOVAs on both the initial MPE (the mean of the first two exposure blocks) and the final exposure MPE (the mean of the last two exposure blocks), treating dwell time as a 3-level factor.

In parallel, we conducted ANOVAs on the mean of the final two exposure blocks for force compensation, with dwell time considered as a 3-level factor. Post-hoc comparisons were conducted using the Holm-Bonferroni correction whenever a significant main effect was identified.

To compare results across experiments, we performed ANOVAs on the force compensation data, specifically using the mean of the final 10 exposure blocks. In this analysis, both experiment condition (2 levels) and dwell time (3 levels) were considered as factors. Again, post-hoc comparisons were conducted with the Holm-Bonferroni correction if a significant main effect was detected.

## Results

Participants performed one of two experiments, one with a visual contextual lead-in, and the other with a passive contextual lead-in, to examine the effects of lead-in modality on adaptation to two opposing force fields. In both experiments, participants first experienced a contextual movement (visual or passive) to a central via-point. After a specified dwell time, they were required to perform an active movement to a given target location.

In each experiment, there were three conditions with different dwell time requirements, which were performed by different groups of participants. These permitted the examination of how adaptation to two opposing curl-fields is affected as the time delay between the contextual movement and the adaptation movement was changed.

Each experiment consisted of a baseline, in which the active movement took place in the null-field condition to acclimatize participants to the experiments and to quantify their baseline movement performance, followed by the introduction of curl force fields introduced to examine adaptation to these novel dynamics (exposure phase). Finally, there was a washout phase.

### Visual Lead-in Experiment 1

#### Dwell Time Analysis

The dwell times correspond to the measured times between the end of the contextual movements and the initiation of the corresponding adaptation movements. To quantify the dwell times that occurred during the training exposure for all participants, we generated a histogram of all curl field trials. This analysis was performed separately for the short, medium, and long experimental conditions. For each histogram, the mean and standard deviation of the dwell time values were calculated to further quantify them.

To quantify the dwell times in trials used to estimate the final levels of compensation achieved by the participants, we also generated a histogram for the final 10 blocks (20 channel trials) in the exposure phase. This analysis was again performed separately for the short, medium, and long experimental conditions.

Fig. 4A shows the histogram plots for the three different dwell time conditions for all exposure trials, and Fig. 4B shows them for the last 10 blocks of channel trials in the exposure phase. We note that there is good agreement in the form of the histograms in the exposure phase and the probe channels used to estimate final levels of compensation, although there are clearly considerably fewer data points in the channel conditions.

**Figure 4.**
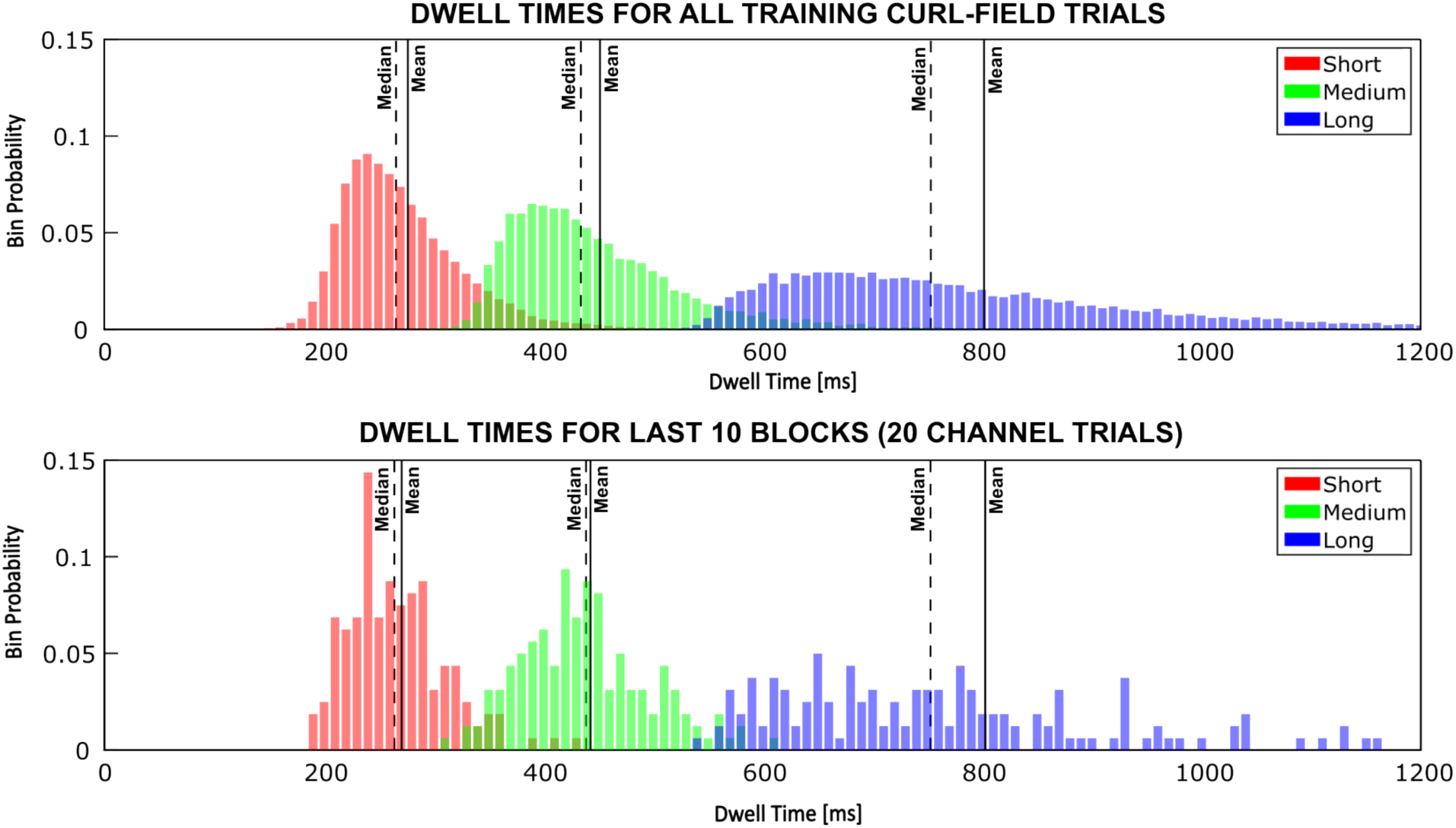
Visual Lead-in Dwell Time Histogram Plots. The plots show the distribution of measured dwell times executed by participants across the three experimental dwell time conditions. The black line indicates the mean dwell time for each condition, and the black dotted line indicates the corresponding median dwell time. **A:** The dwell time histogram for curl field exposure trials shows mean (median) values of 276.9 ms (266.0 ms) for short, 452.1 ms (434.6 ms) for medium, and 802.2 ms (753.5 ms) for long dwell times. **B:** The dwell time histogram for the channel trials during the last 10 blocks (20 channel trials) of the exposure phase shows mean (median) values of 270.7 ms (264.3 ms) for short, 442.9 ms (439.0 ms) for medium, and 802.6 ms (753.0 ms) for long dwell times.

In addition, in all cases, the mean and mode values differ from each other, and the mode of the distributions is shifted from the mean to the left, towards lower values. The dwell time distributions also become more spread out for the larger dwell time conditions.

### Hand Trajectories

The hand trajectories for all three conditions in the Visual Lead-in Experiment 1 were examined in four different stages of the experiments (Fig. 5). The first column (Last Null) shows hand trajectories during the last block of trials in the null phase and shows that participants were able to perform unhindered straight movements in all three conditions. The second column (Initial Curl) shows the first block of trials after the introduction of the curl-field, revealing that participants deviated strongly from a straight movement in all three conditions. The third column (Final Curl) shows hand trajectories during the last block of curl-field trials. This highlights distinct differences across dwell time conditions. Notably, in the short dwell time condition, participants performed almost straight movements, close to baseline performance. In the medium dwell time condition, participants still exhibited effects of the curl-fields and still deviated somewhat from the straight path. In the long dwell time condition, the effect of the curl-fields was still apparent, suggesting there was little adaptation. The final column (Washout) shows the first block of trials during the washout phase, after the curl-field was removed. It is evident that short dwell times led to strong aftereffects, leading to significant deviation from straight trajectories in the opposite direction to that seen during curl-field exposure. As the dwell time increased, the deviation was reduced, and in the long dwell time condition, the trajectories were effectively straight.

**Figure 5.**
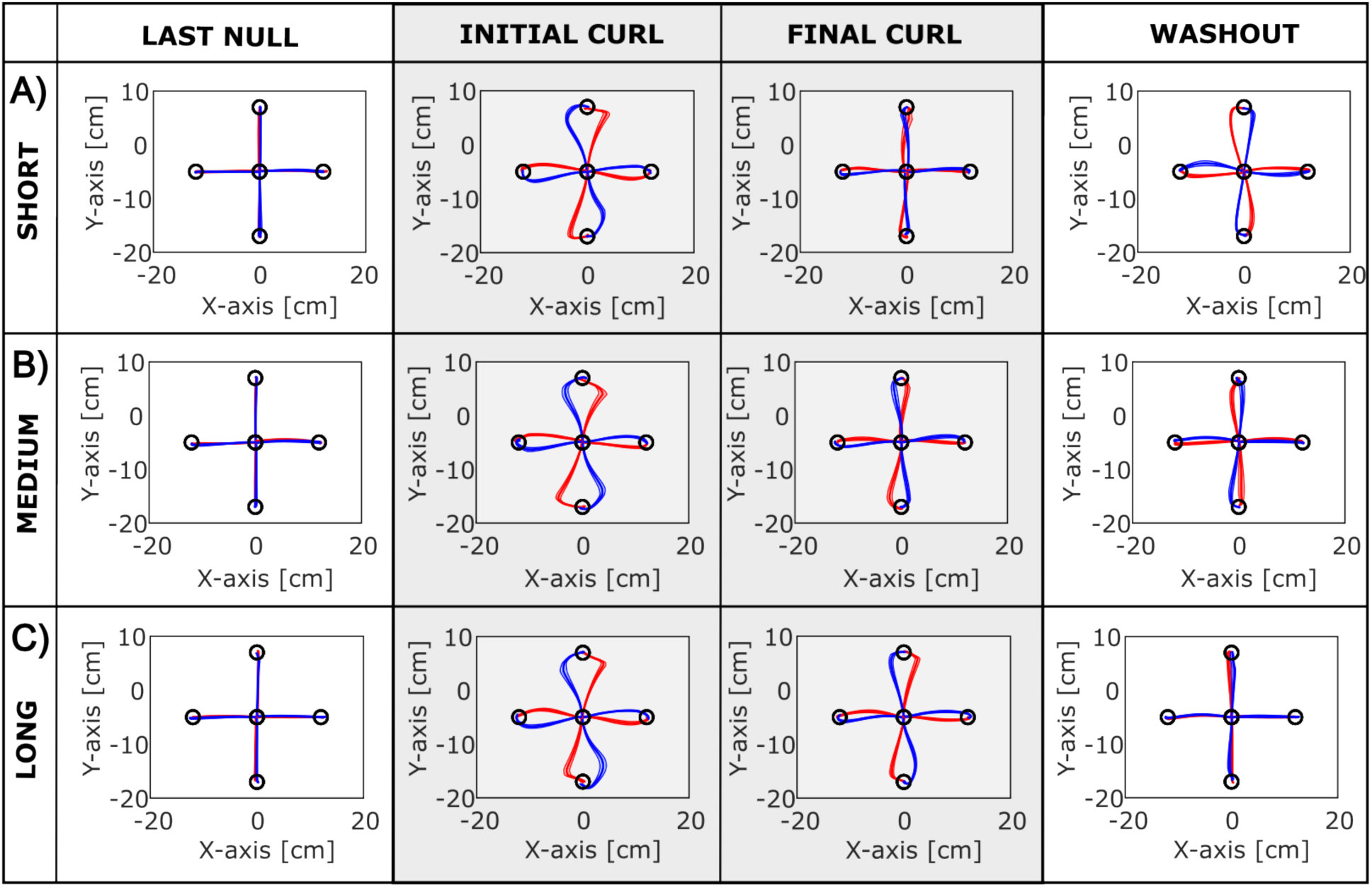
Hand Trajectories in the Visual Lead-in Experiment 1. The solid thick lines indicate the mean across participants and the shaded region indicates the standard error of the mean. Red colors indicate trajectories in the context associated with CW curl force fields whereas blue colors indicate the trajectories in the context associated with CCW curl force fields. **A:** The short dwell time condition. **B:** The medium dwell time condition. **C:** The long dwell time condition. The columns represent the Last Null, Initial Curl, Final Curl, and Washout stages of the experiment.

### Kinematic Error and Force Compensation

To examine the overall level of adaptation in the different dwell conditions, we initially combined the effect of both curl-field directions by calculating a single value from the dual Maximum Perpendicular Error (MPE) and compensation values (Fig. 6).

**Figure 6.**
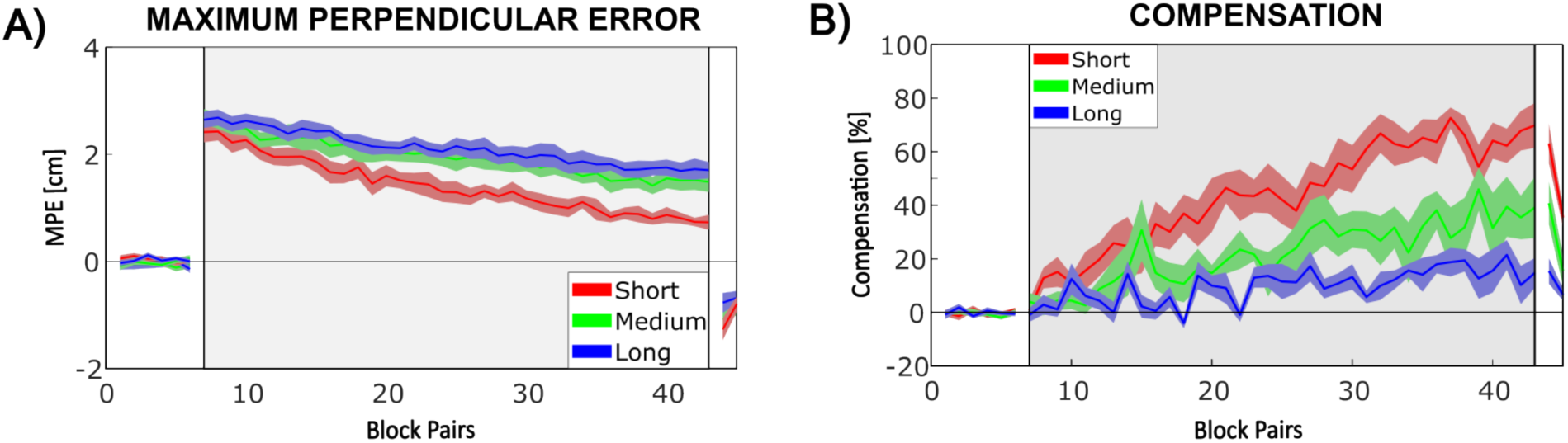
Adaptation in the Visual Lead-In Experiment 1. **A:** Kinematic error. Overall Mean (solid line) ± SE (shaded region) of the MPE plotted against block pairs for the short, medium, and long dwell-time conditions. **B:** Force Compensation. Overall Mean ± SE percentage of compensation for the short, medium, and long dwell-time conditions.

To examine kinematic error during the experiments we plotted the Maximum Perpendicular Error (MPE) across the experimental blocks for the short, medium, and long dwell time conditions (Fig. 6A). During the null-field exposure, MPE was essentially zero across all conditions. Upon the introduction of the curl-field, MPE dramatically increases in all three conditions. However, by the end of curl-field exposure, MPE decreases by a larger amount in the case of short dwell times compared to long dwell times. In this case, MPE for the medium dwell time was only slightly lower than for the longer dwell times. During the final washout phase, the MPE exhibited a negative sign, indicating deviation in the opposite direction of that created by the curl-field. Short dwell times led to a larger magnitude of after effect than medium or long dwell times.

Repeated measures ANOVAs indicated that the MPE showed significant variations between the pre-exposure, initial exposure, final exposure, and washout phases for all dwell time conditions. Post-hoc comparisons further tested the significant differences between these phases. In the short dwell time condition (F₃,₂₁ = 121.748; p < 0.001; ω² = 0.903), MPE was low during the initial null-field exposure, then it significantly increased upon introduction of the curl-field (p_bonf_ < 0.001). Over the exposure phase, MPE significantly decreased (p_bonf_ < 0.001). MPE during the washout phase showed a significant difference compared to the pre-exposure phase (p_bonf_ < 0.001). For the medium dwell time condition (F₃,₂₁ = 176.789; p < 0.001; ω² = 0.929), MPE was low during the initial null-field exposure, and MPE increased significantly upon the introduction of the curl-field (p_bonf_ < 0.001). Over the exposure phase, MPE decreased significantly (p_bonf_ < 0.001). MPE during the washout phase showed a significant difference compared to the pre-exposure phase (p_bonf_ = 0.007). In the long dwell time condition (F₃,₂₁ = 220.335; p < 0.001; ω² = 0.926), MPE increased significantly upon the introduction of the curl-field (p_bonf_ < 0.001). Over the exposure phase, MPE decreased significantly (p_bonf_ < 0.001). However, MPE during the washout phase showed no significant difference compared to the pre-exposure phase (p_bonf_ = 0.669).

To quantify the amount of predictive force compensation to the force fields, we examined the force compensation on the channel trials for the short, medium and long conditions (Fig. 6B). During the initial null-field exposure, compensation was low in all dwell time conditions. Upon the introduction of the curl-field, compensation gradually increased throughout the exposure period. By the end of the curl-field exposure, the compensation for the short dwell time condition increased the most, while for longer dwell times, the least compensation was observed. An intermediate compensation level was seen for the medium dwell time. During the washout phase, compensation values decayed in all three dwell time conditions. Repeated measures ANOVAs indicated a significant increase in force compensation between the final pre-exposure and final exposure levels for short dwell times (F_1,7_ = 59.253; p < 0.001; ω² = 0.785), medium dwell times (F_1,7_ = 28.357; p = 0.001; ω² = 0.617), and long dwell times (F_1,7_ = 8.366; p = 0.023; ω² = 0.280).

The MPE and compensation results indicate that participants were better able to compensate opposing dynamics by making use of the visual lead-in context at shorter dwell times. The contextual effect decreased as the dwell time increased. Observations from the trajectory plots (Fig. 5) are consistent with these findings.

### Independent Adaptation to Opposing Force Fields

While the previous analysis clearly shows adaptation to the dynamics, it is unclear the level to which this is independently adapted to each force field direction. Fig. 7 shows the Maximum Perpendicular Error (MPE) and compensation for the Visual Lead-In Experiment 1, with both plotted separately for the two contextual directions, to permit the individual examination of the learning process in each curl-field direction. Similar trends are observed in both curl-field directions.

**Figure 7.**
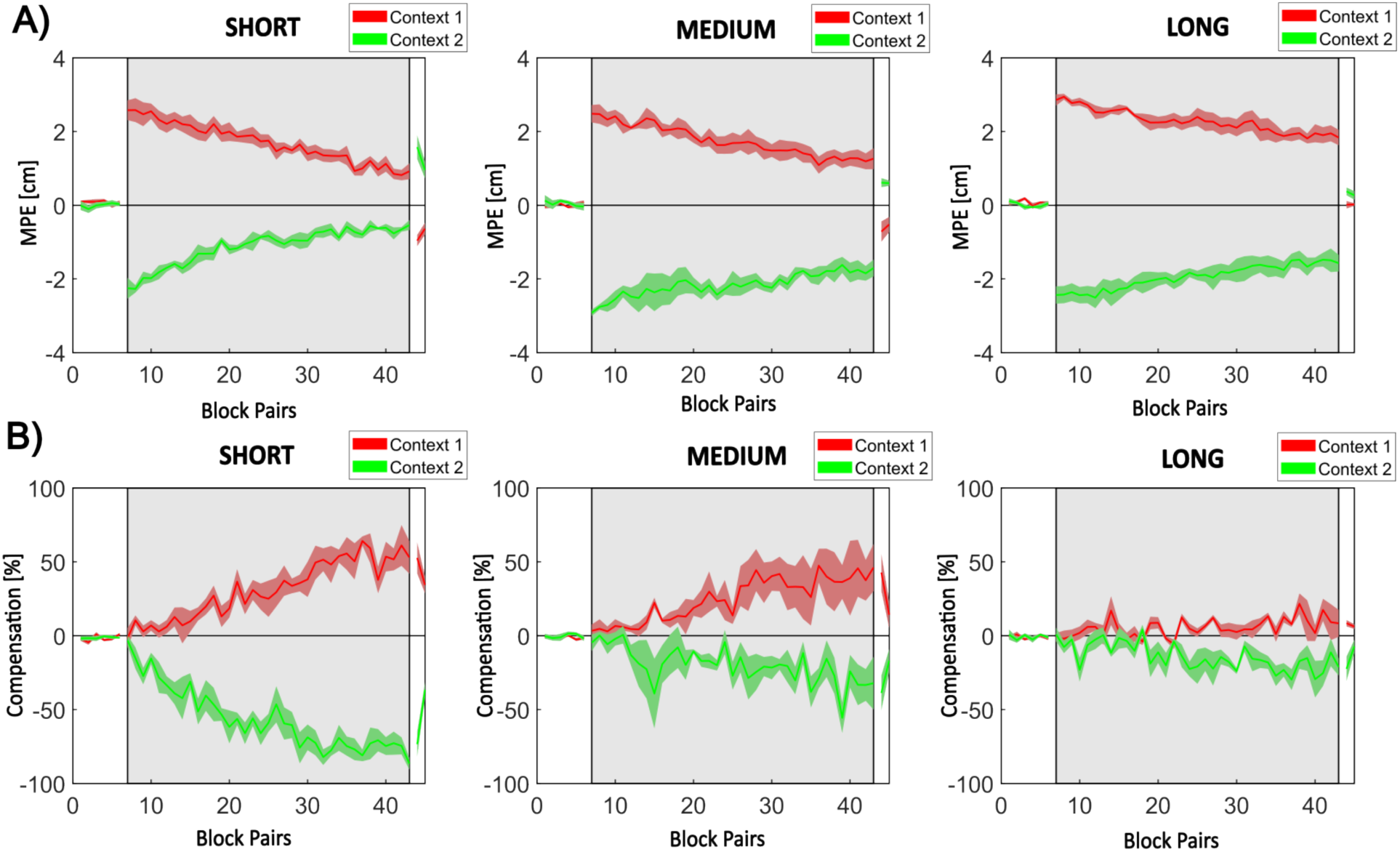
Independent Adaptation to Opposing Force Fields. Visual Lead-In Mean ± SE Maximum Perpendicular Error (MPE) and Compensation are plotted for both opposing curl-fields. **A:** Mean ± SE MPE for short, medium, and long dwell times. **B:** Mean ± SE compensation for short, medium, and long dwell times.

We compared the final exposure MPE in both curl-field directions of block pairs across participants (Fig. 7A). A repeated measures ANOVA indicated that there were no significant differences between the two directions for the short dwell time condition (F_1,7_ = 1.199; p = 0.310; ω² = 0.004), for the medium dwell time condition (F_1,7_ = 0.246; p = 0. 645; ω² = 0.000), or for the long dwell time condition (F_1,7_ = 3.501; p = 0.104; ω² = 0.034).

Similarly, Fig. 7B shows the mean ± SE compensation for short, medium, and long dwell times. We compared the final exposure compensation in both curl-field directions of block pairs across participants. A repeated measures ANOVA indicated that they were not significantly different, for the short dwell time condition (F_1,7_ = 0.023; p = 0.883; ω² = 0.000), for the medium dwell time condition (F_1,7_ = 0.131; p = 0.728; ω² = 0.000), or for the long dwell time condition (F_1,7_ = 3.501; p = 0.433; ω² = 0.000).

### Passive Lead-in Experiment 2

Dwell Time Analysis: Following a similar analysis for the visual lead-in data, Fig. 8A shows the histogram plots for the three different dwell time conditions for all exposure trials, and Fig. 8B shows them for the final 10 blocks (20 channel trials) in the exposure phase. We note that there is reasonable agreement in the form of the histograms in the exposure phase and the probe channels used to estimate final levels of compensation at lower dwell times, but there appears to be a shift in the distribution for channel trials towards lower dwell times in the long condition.

**Figure 8.**
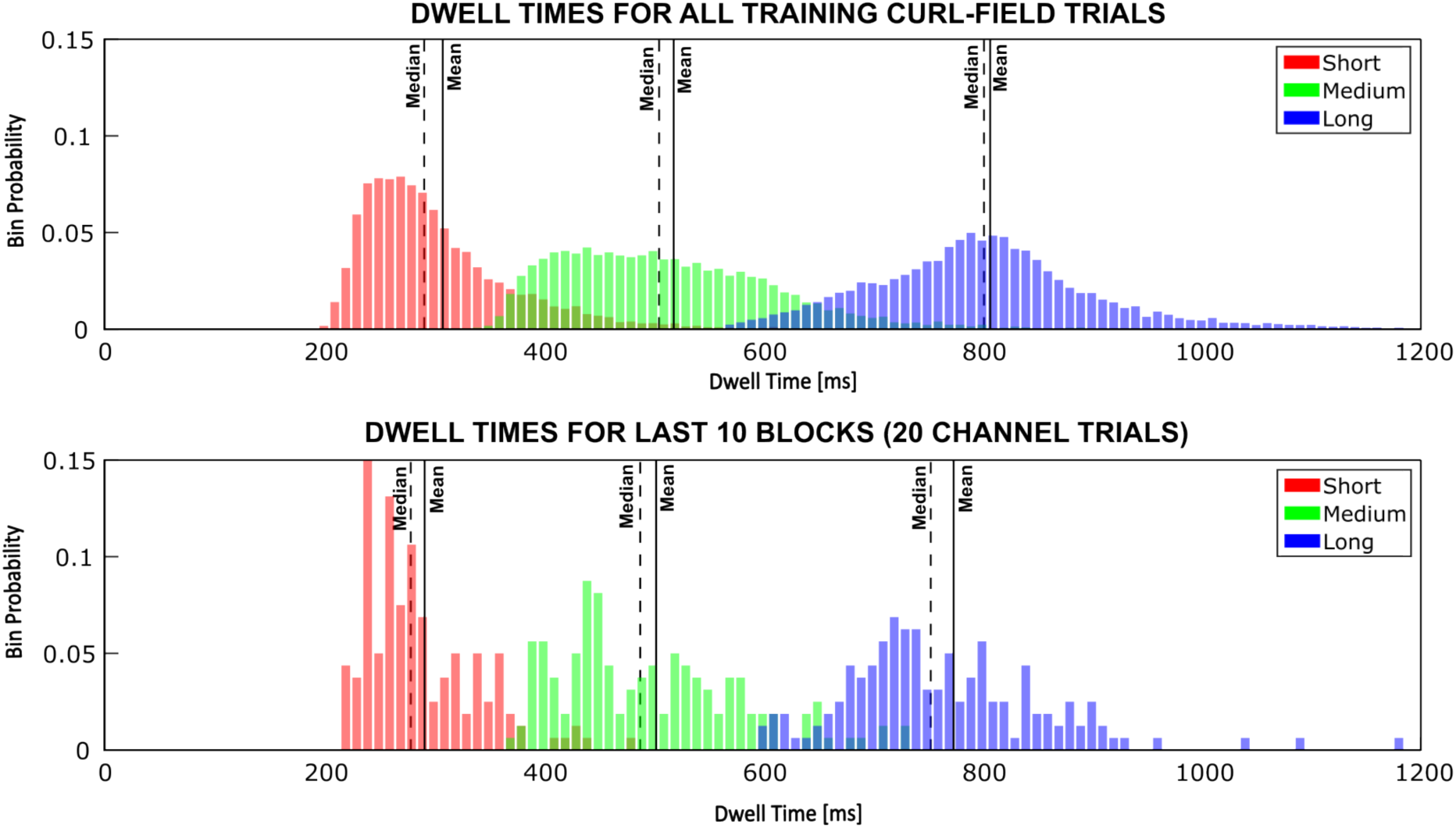
Passive Lead-in Dwell Time Histogram Plots. The histograms show the distribution of measured movement dwell times in the lead-in movements executed by participants across the three experimental dwell time conditions. The black line indicates the mean dwell time for each condition, and the black dotted line indicates the corresponding median dwell time. **A:** The dwell time histogram for curl field exposure trials shows mean (median) values of 308.5 ms (291.3 ms) for short, 519.0 ms (505.4 ms) for medium, and 807.5 ms (801.6 ms) for long dwell times. **B:** The dwell time histogram for the channel trials during the last 10 blocks (20 channel trials) of the exposure phase shows mean (median) values of 291.9 ms (280.0 ms) for short, 503.0 ms (489.1 ms) for medium, and 774.1 ms (754.0 ms) for long dwell times.

The distributions spread out slightly as dwell time increases. In the short condition, the mean and mode values deviate and are shifted to the left of the mean. However, in the long condition, the mode and mean times are much closer, and the distributions become more symmetrical around the mean.

### Hand Trajectories

Hand trajectories were computed across the four stages for each of the three dwell time conditions (Fig. 9). As shown in the first column (Last Null), participants were able to perform unhindered straight movements in all three dwell time conditions. The second column (Initial Curl) shows that on initial curl-field exposure, participants deviated strongly from a straight movement in all three conditions. The third column (Final Curl) shows that by the end of the training phase, participants’ hand movements were almost straight in the short dwell time condition but deviated more from a straight line as the dwell time increased. However, even in the long dwell time condition, they were still straighter than those seen in the initial exposure phase. The final column (Washout) shows strong aftereffects for the short dwell time condition. As the dwell time increases, aftereffects become less prominent although there is still some deviation from a straight-line trajectory even in the long dwell time condition.

**Figure 9.**
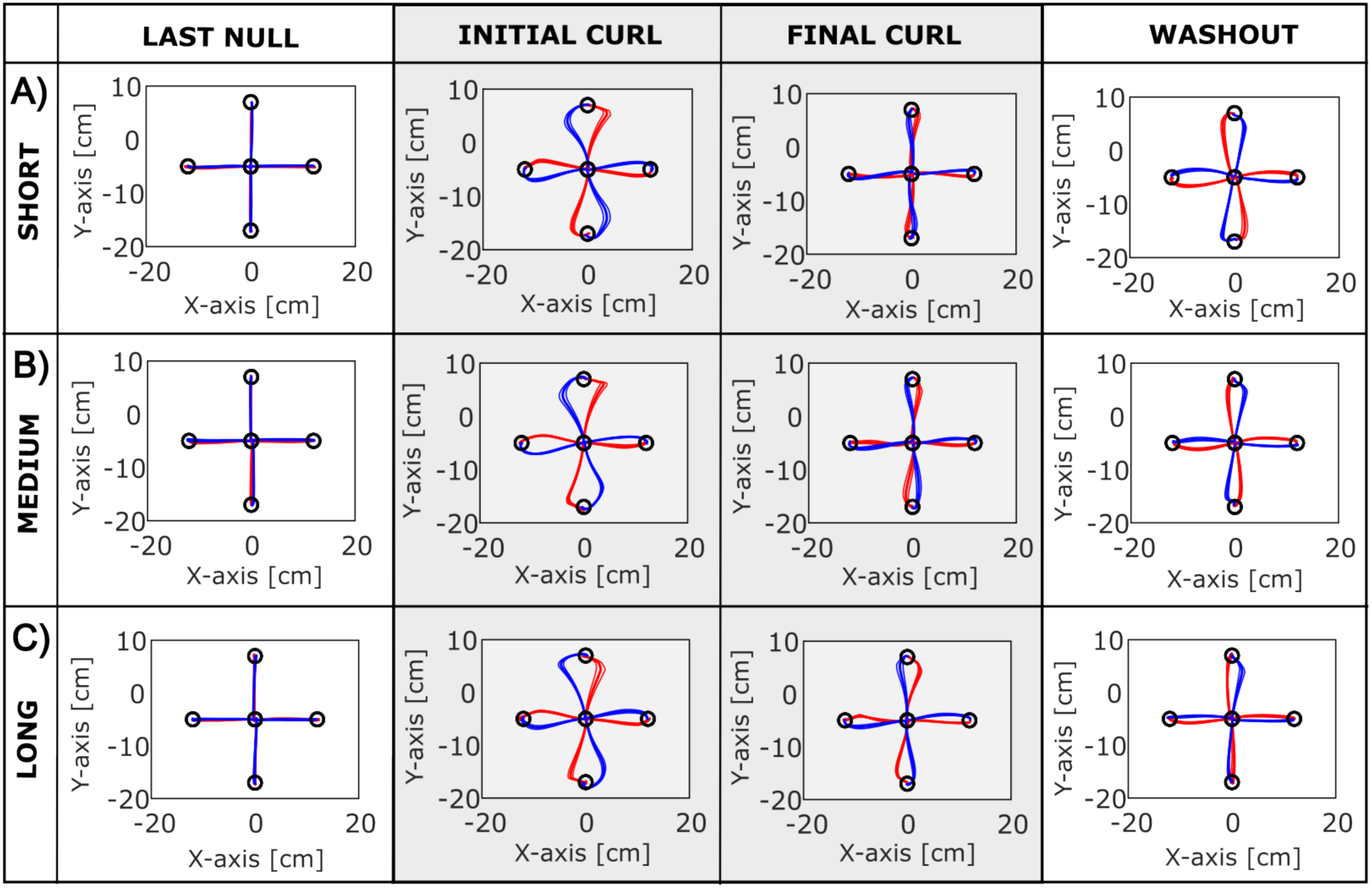
Hand Trajectories for the Passive Lead-In Experiment 2. Plot of the Mean ± SE Hand Trajectories for the three experiment conditions. **A:** The short dwell time condition. **B:** The medium dwell time condition. **C:** The long dwell time condition. The columns represent the Last Null, Initial Curl, Final Curl, and Washout stages of the experiment.

### Kinematic Error and Force Compensation

To combine the effect of both curl-field directions, we calculated a single value from the dual Maximum Perpendicular Error (MPE) and compensation values (Fig. 10). During the null-field exposure, MPE was essentially zero across all conditions (Fig. 10A). Upon the introduction of the curl-field, MPE dramatically increased in all three conditions. By the end of curl-field exposure, MPE decreased more with shorter dwell times than with longer dwell times. During the washout phase, the MPE exhibited a negative value.

**Figure 10.**
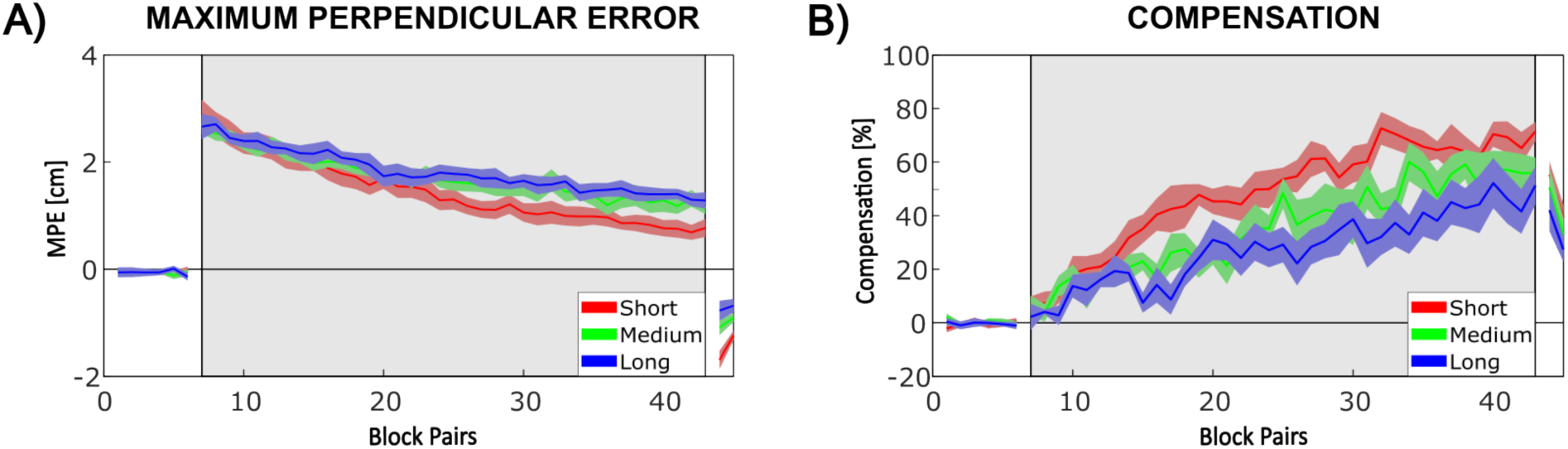
MPE and Compensation for the Passive Lead-in Experiment 2. **A:** Overall Mean ± SE MPE plotted against block pairs for each of the three dwell-time conditions. **B:** Overall Mean±SE percentage of compensation for each of the three dwell-time conditions.

Repeated measures ANOVAs revealed significant differences in MPE across the pre-exposure, initial exposure, final exposure, and washout phases for all dwell time conditions. In the short dwell time condition, significant variations were observed (F_3,21_ = 134.537, p < 0.001, ω² = 0.909). Post-hoc comparisons showed that MPE was low during the initial null-field exposure but increased significantly upon the introduction of the curl-field (p_bonf_ < 0.001). Over the exposure phase, MPE decreased significantly (p_bonf_ < 0.001), and during the washout phase, MPE showed a significant difference compared to pre-exposure (p_bonf_ < 0.001). In the medium dwell time condition, MPE also showed significant differences across the phases (F_3,21_ = 186.893, p < 0.001, ω² = 0.924). During the initial null-field exposure, MPE was low, but it increased significantly with the introduction of the curl-field (p_bonf_ < 0.001). MPE decreased significantly over the exposure phase (p_bonf_ < 0.001), and in the washout phase, MPE remained significantly different from the pre-exposure phase (p_bonf_ = 0.002). For the long dwell time condition, significant variations were found (F_3,21_ = 101.650, p < 0.001, ω² = 0.889). MPE was initially low during the null-field exposure and increased significantly with the curl-field introduction (p_bonf_ < 0.001). Over the exposure phase, MPE decreased significantly (p_bonf_ = 0.002), and the washout phase showed a significant difference in MPE compared to the pre-exposure phase (p_bonf_ = 0.026).

The mean force compensation was examined (Fig. 10B) for the short, medium, and long dwell time conditions. During the initial null-field exposure, compensation was low in all dwell time conditions. Upon the introduction of the curl-field, compensation began to increase. By the end of the curl-field exposure, the compensation had increased for all dwell time conditions, but less so as dwell time increased. During the washout phase, compensation values decayed in all three dwell time conditions.

A repeated measures ANOVA indicated a significant increase in force compensation between the final pre-exposure and final exposure levels across all dwell time conditions. In the short dwell time condition, the increase was highly significant (F_1,7_ = 182.743, p < 0.001, ω² = 0.929). For the medium dwell time condition, a significant increase in force compensation was also observed (F_1,7_ = 30.547, p < 0.001, ω² = 0.664). Similarly, in the long dwell time condition, a significant increase in force compensation between the final pre-exposure and final exposure levels was found (F_1,7_ = 39.970, p < 0.001, ω² = 0.693).

These MPE and force compensation results demonstrate that participants using passive lead-in contexts could effectively compensate for both opposing curl-field directions, consistent with the raw trajectory plots shown in Fig. 9. Although the contextual effect diminished as the dwell time increased, it decreased less than what was observed in Visual Lead-in Experiment 1.

### Independent Adaptation to Opposing Force Fields

The Maximum Perpendicular Error (MPE) and force compensation were examined for each of the two curl force field directions for the Passive Lead-In Experiment 2 to test if there are differences in the adaptation of the two field directions (Fig. 11).

**Figure 11.**
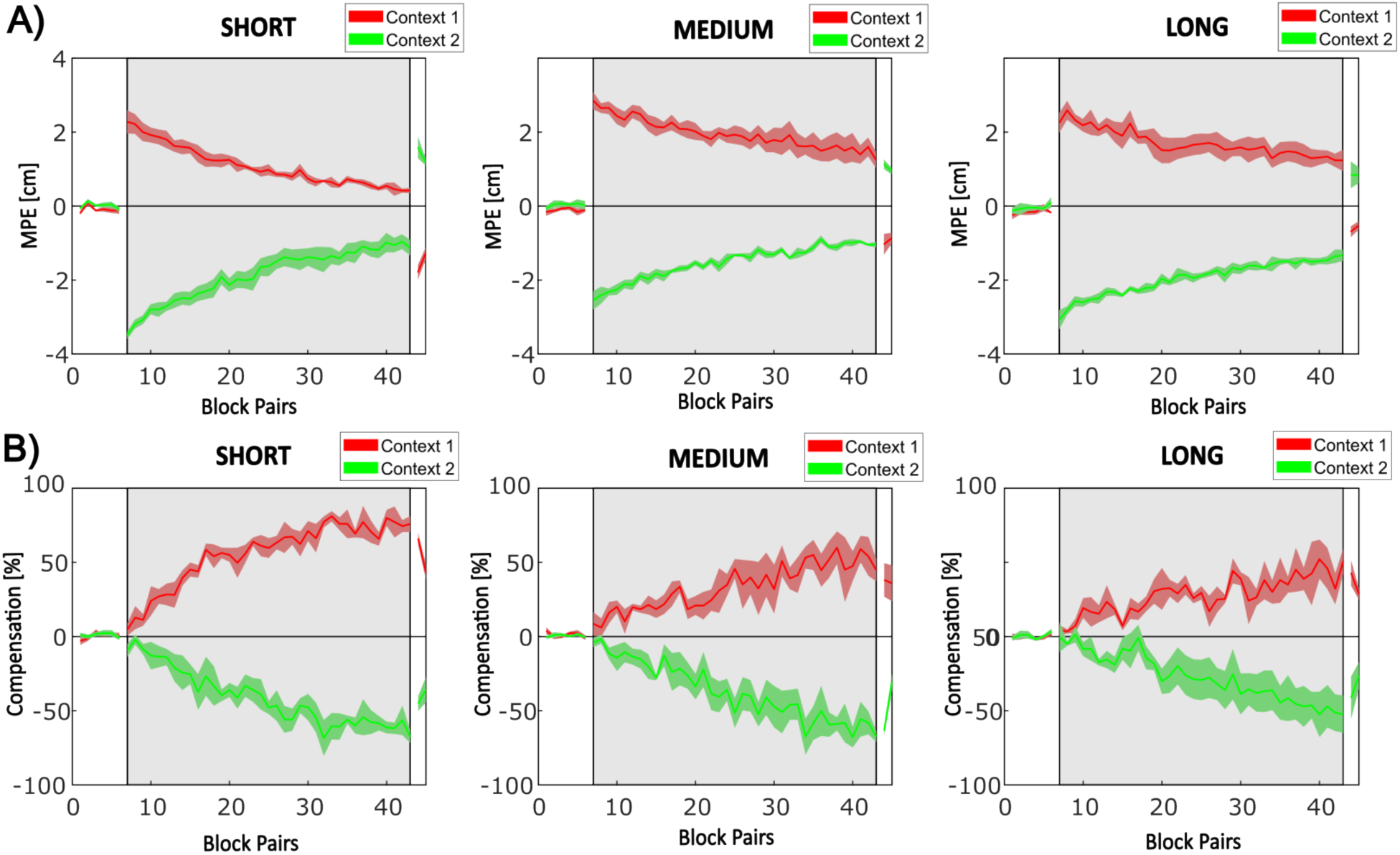
Independent Adaptation to Opposing Force Fields for the Passive Lead-In. The mean ± SE MPE and force compensation for both opposing curl-fields. **A:** Mean ± SE MPE for short, medium, and long dwell times. **B:** Mean ± SE compensation for short, medium, and long dwell times.

Similar trends in the MPE are seen in both curl-field directions (Fig. 11A). We compared the final exposure MPE across participants for both curl-field directions within block pairs. A repeated measures ANOVA indicated that there were no significant differences in adaptation across curl-field directions for the short, medium, and long dwell time conditions. Their analyses yielded the following statistics: (F_1,7_ = 0.231; p = 0.645; ω² = 0.000), (F_1,7_ = 3.543; p = 0. 102; ω² = 0.072), and (F_1,7_ = 2.178; p = 0. 184; ω² = 0.059) respectively. That is, participants adapted equally to both force field directions.

We compared the final exposure compensation (Fig. 11B) in both curl-field directions of block pairs across participants. A repeated measures ANOVA indicated that there were no significant differences for the medium and long dwell time conditions F_1,7_ = 0.268; p = 0. 621; ω² = 0.000) and (F_1,7_ = 2.780; p = 0.139; ω² = 0.067) respectively). However, a significant difference was observed in the short dwell time condition (F_1,7_ = 6.078; p = 0.043; ω² = 0.210).

### Comparison of Results Across Experimental Conditions

#### Adaptation during Curl Field Exposure

To contrast adaptation across the two experiments, we performed comparisons of the levels of curl-field compensation achieved in both the Visual Lead-in Experiment 1 and Passive Lead-in Experiment 2 (Fig. 12). At short dwell times (Fig. 12A), compensation levels increased similarly in both experiments during curl-field exposure. At medium dwell times (Fig. 12B), a slight divergence emerged, with the Visual Lead-in Experiment 1 showing a slower increase in learning compared to the Passive Lead-in Experiment 2. This difference became more pronounced at long dwell times (Fig.12C), where the Visual Lead-in Experiment 1 exhibited noticeably less learning than the Passive Lead-in Experiment 2.

**Figure 12.**
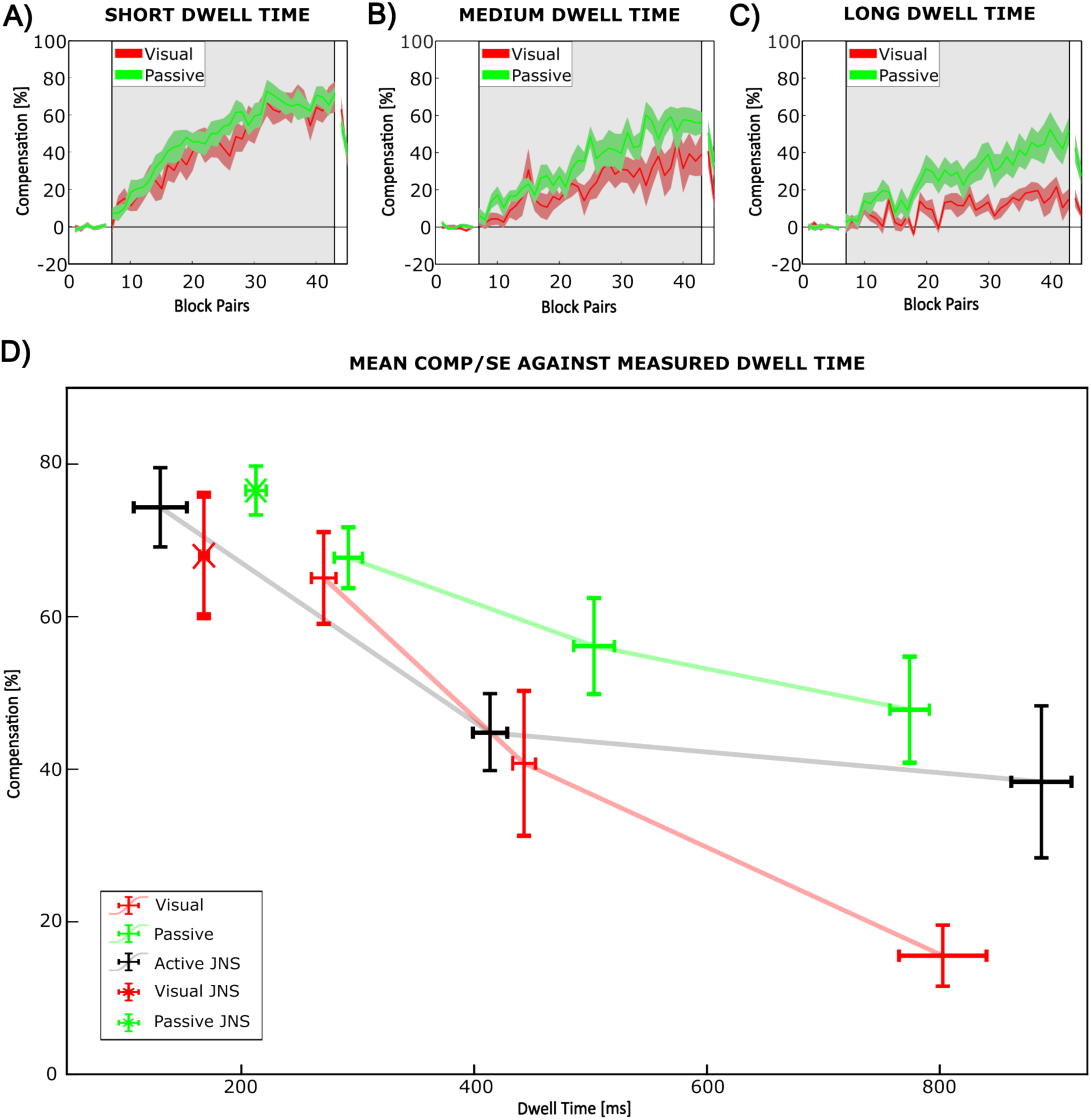
Comparison of adaptation across the experiments. Plot of mean±SE compensation values for the Visual Lead-in Experiment 1 and the Passive Lead-in Experiment 2 contexts across the three different dwell time conditions. **A, B & C:** Force compensation against block pairs for both experiments at short, medium and long dwell times, respectively. **D:** Final level of curl-field compensation (mean±SE) plotted against the measured dwell time (mean±SE). This includes results from both experimental conditions in the current study, as well as results from a previous study that examined an active lead-in context (15). In addition, the plot includes low dwell time results for visual and passive lead-ins from the previous study.

An ANOVA analysis of the last 10 curl-field exposure trial blocks was performed for both the Visual Lead-In Experiment 1 and the Passive Lead-In Experiment 2. Final exposure compensation showed significant variations between dwell-time conditions (F₂ = 14.685; p < 0.001; ω² = 0.311) and experimental conditions (F₁ = 10.257; p =0.003; ω² = 0.105). Post-hoc comparisons revealed no significant differences between Visual Lead-in Experiment 1 and Passive Lead-in Experiment 2 for short and medium dwell times (p_bonf_ =1.000). However, there was a significant difference for long dwell times (p_bonf_ =0.014).

We note that, even though the passive modality generally exhibited longer mean dwell time values than the visual modality, we still saw significantly greater adaptation in the Passive Lead-in Experiment 2 than in the Visual Lead-in Experiment 1. This indicates that it is the form of the lead-in modality that is responsible for the observed differences in adaptation achieved since the Passive Lead-in Experiment 2 showed greater adaptation even though dwell times were slightly longer in the passive condition compared to the visual condition. These findings indicate a more rapid decay of contextual effectiveness when using visual lead-in cues compared to passive lead-ins. Furthermore, the data suggest that the effectiveness of visual lead-in cues substantially declines when dwell times exceed approximately 800 ms.

#### Comparison with Previous Studies

To further examine the relationship between lead-in modality and the decay of compensation as a function of dwell time across different experimental conditions, the mean compensation values calculated in channel trials over the last 10 blocks of curl-field exposure trials were plotted against the mean of their measured dwell times.

Fig. 4 and Fig. 8 show that the histograms of dwell times for the different experimental conditions do not follow Gaussian distributions; indeed, some exhibit substantial asymmetry. Therefore, the mean and standard deviation do not constitute sufficient statistics to represent dwell time distributions. However, we note from observing the histograms that the mean and median values do not substantially differ, although the mode values, representing the peak each histogram, are certainly shorter. For simplicity, we chose the mean dwell time as a representative metric for plotting.

As well as performing this analysis on the passive and visual lead-in data collected in the current study, we also relate our new results to previous work that examined the effect of dwell time on an active lead-in context, which also reported compensation values achieved with short dwell times for passive and visual lead-in contexts (15). To do so, we fully re-analyzed the appropriate raw data generated in the former study. A detailed breakdown of the compensation, MPE, and dwell time values for the last 10 blocks of exposure trials for all current experimental conditions and relevant previous experimental conditions is provided in Table 2. To facilitate comparison between the current data and reanalysed data, dwell time conditions were grouped into four bins based on their durations: Very Short (“V-Short”), Short, Medium, and Long. This arbitrary bin selection was intended only to broadly classify dwell times, and since not all conditions arose in both sets of experiments, the tables are consequently not fully populated.

**Table 2.** Measured mean and standard error or dwell times and compensation values over the last 10 blocks of channel trials in the exposure phase for the visual, passive and active lead-In conditions (mean±SE). The standard deviation (Std) over all the trials is also provided. Results from the former study are shaded in grey.

|  | Visual Lead-In |  | Passive Lead-in |  | Active Lead-in |  |
| --- | --- | --- | --- | --- | --- | --- |
|  | COMP<br>mean±SE<br>Std | DWELL<br>mean±SE<br>Std | COMP<br>(mean±SE<br>Std | DWELL<br>(mean±SE<br>Std | COMP<br>(mean±SE<br>Std | DWELL<br>(mean±SE<br>Std |
| **V-Short** | 68.0 ± 8.0% | 167.8 ± 3.1<br>ms | 76.5 ± 3.2% | 212.5 ± 8.7<br>ms | 74.3 ± 5.2%<br>23.2% | 130.3 ± 22.9<br>ms<br>69.5 ms |
| **Short** | 65.1 ± 6.0%<br>30.8% | 270.7 ± 10.7<br>ms<br>56.2 ms | 67.7 ± 4.0%<br>22.5% | 291.9 ± 12.2 ms<br>72.0ms | - | - |
| **Medium** | 40.8 ± 9.5%<br>43.0% | 442.9 ± 8.4<br>ms | 56.2 ± 6.3%<br>32.3% | 503.0 ± 17.5 ms | 44.785 ± 5.1214% | 413.6 ± 14.8<br>ms |
|  |  | 82.2 ms |  | 100.3 ms | 35.141% | 83.6 ms |
| **Long** | 15.6 ± 4.1%<br>35.2% | 802.6 ± 37.6 ms<br>200.3 ms | 47.8 ± 7.0%<br>40.6% | 774.15 ± 16.9 ms<br>100.5 ms | 38.4 ± 10.0%<br>34.8% | 887.3 ± 25.9 ms<br>137.1 ms |

Although the mean duration of the lead-in movements differs across the visual, passive, and active experiments (see Table 3), these differences did not substantially affect the level of adaptation achieved at short dwell times across the experimental conditions, since the training and testing dwell times were similar.

**Table 3.** Measured lead-in movement durations over the last 10 blocks of exposure trials for the visual, passive and active lead-In conditions (mean±SE). Results from the former study are shaded in grey. We note that the SE and SD values for the visual and passive lead-in conditions are small because they are highly consistent, since they are generated by the robotic manipulandum system.

|  | Visual Lead-In | Passive Lead-in | Active Lead-in |
| --- | --- | --- | --- |
|  | Lead-In Duration<br>mean±SE<br>Std | Lead-In Duration<br>mean±SE<br>Std | Lead-In Duration<br>mean±SE<br>Std |
| **V-Short** | 551.1 ± 18.1 ms<br>48.1 ms | 582.0 ± 23.7 ms<br>54.5 ms | 204.0 ± 12.4 ms<br>43.8 ms |
| **Short** | 514.7 ± 0.1 ms<br>0.7 ms | 571.1 ± 1.8 ms<br>15.1 ms | - |
| **Medium** | 514.6 ± 0.1 ms<br>0.7 ms | 570.8 ± 1.9 ms<br>15.41 ms | 224.8 ± 7.3 ms<br>33.0 ms |
| **Long** | 514.7 ± 0.1 ms<br>0.7 ms | 571.0 ± 2.6 ms<br>16.6 ms | 230.7 ± 12.3 ms<br>49.0 ms |

Fig. 12D highlights a clear contrast between the visual, passive and active lead-in experiments. In the Visual Lead-in Experiment 1, compensation declined markedly as dwell time increased. In contrast, although the Passive Lead-in Experiment 2 also showed a decline in compensation with increasing dwell time, the slope was shallower (i.e., less negative), and overall compensation levels remained consistently higher than those observed in the visual condition. After reanalysis, the previously collected active data exhibit a decline in force compensation with increasing dwell time, falling between the compensation values observed in the new passive and visual lead-in conditions.

The reanalysis of the low-dwell passive and visual conditions from the former study (also shown in Fig. 12D as individual data points) aligns well with the trends seen in the new passive and visual data, constituting data points with higher compensation values at lower dwell times.

## Discussion

### Summary

Using an interference paradigm, we investigated the contextual effects of visual and passive lead-in movements on motor adaptation as a function of dwell time. Our results revealed significant differences between the two experimental conditions. We found that in both lead-in modalities, the level of adaptation achieved decreased as dwell time increased. However, visual lead-in movements showed a greater decline compared to passive lead-in movements. Although adaptation in both visual and passive lead-in conditions was similar at shorter dwell times, the level of adaptation achieved with visual lead-ins became much smaller when measured dwell time exceeded approximately 800 ms. In contrast, significantly more adaptation was present in the passive lead-in condition.

### Comparison with a former Lead-In Study

Our current study closely followed the experimental protocol of the previous active lead-in movement study on the effect of dwell time (15), but we refined the analysis methods. Specifically, we examined the final levels of compensation using the last 10 blocks of field exposure rather than the last 25, providing a more precise estimate of the final adaptation achieved in each experimental condition. Additionally, we calculated compensation by regressing movement velocity against measured force instead of using the ratio of velocity to force, and we implemented a more robust method to estimate dwell time. To enable meaningful comparisons with previous results, we reanalyzed the relevant data from the earlier study using these updated methods.

Firstly, we observed that compensation values for passive and visual lead-ins from the former study under low dwell time conditions closely matched the trends seen in our visual and passive data (see Fig. 12). Furthermore, our new results align with previous findings on active lead-in contextual movements, which demonstrated a decline in the contextual effect and a subsequent reduction in adaptation as dwell time increased (15). We note that compensation levels in the active lead-in condition were slightly lower than those in the passive condition, but higher than in the visual condition. In the former case, the key factor affecting learning may arise from the higher levels of lead-in variability in the active condition compared to the passive lead-in condition. Previous studies have shown that increased variability reduces adaptation (23), which suggests that the lower compensation observed with active lead-ins may be due to the higher variability inherent in self-generated movements compared to the consistent, system-generated passive movements. The fact that both passive and active lead-in movements achieve high levels of compensation suggests that visual lead-in is a fundamentally weaker effect.

### Effectiveness of Contextual Cues

A contextual cue is a type of information used by the motor system that can influence motor adaptation but is not directly involved in a movement. It has been proposed (38) that there are two main types of contextual cues: direct and indirect. The indirect cues, such as color, often have little effect on implicit motor adaptation (13, 38). However, direct cues, which might connect to state estimation, have strong contextual effects and drive implicit motor adaptation. Many dynamic learning dual-adaptation interference studies have shown that sensory information related to movement has a strong contextual cue effect. Using different physical workspace locations as a context works effectively, as does simply showing different visual locations in the workspace when the actual movements are always made in the same location (13). Immediate past movement (15) or planned future movement also has a very strong contextual effect (39, 40). However, the actual movement itself needs to be somehow associated with the task, and peripheral movement has a much weaker effect than something that looks like a contiguous movement carried out by the participant. It was also found that past visual or passive lead-in movement strongly affects motor learning in subsequent movements. In contrast, peripheral visual movement has a much weaker effect. Here we show that, similar to actively generated lead-in movements, both passive lead-in and visual lead-in movements produce strong contextual cue effects that decay with increasing dwell times between the lead-in and adaptation movement.

### Visual and Proprioceptive Feedback Pathways

The control of active, visually guided arm movements, including the point-to-point movements examined in this study, generally involve integrating visual and proprioceptive sensory feedback within the brain. Visual information requires complex processing in the visual cortex and higher brain centers, introducing delays and potential degradation over time (41–43). In visually guided movements, vision provides not only the location of the desired target but also an online indication of limb position and its movement (44). Visual feedback is essential for planning and guiding movements over extended time scales. However, even the fastest visual feedback is subject to delays of at least 100 to 150 ms due to the pathways and extensive processing required (45, 46).

In contrast, proprioceptive information, which originates from sensors in the muscle fibers, tendons, and joints, is transmitted much faster (47), resulting in delays of approximately 30 to 50 ms (48). Proprioception provides continuous, lower-latency feedback about the body’s position and movement (47, 49). Such proprioceptive feedback is crucial for rapid, reflexive compensatory movements, and multi-joint control, as is clearly demonstrated in patients who experience deficits in proprioception (50).

Thus, visual feedback has a longer time delay compared to proprioceptive feedback, primarily due to the extended time required to process signals, especially for pathways which go through the visual cortex and their subsequent integration into higher brain centers (51). In general, different sensory feedback delays can range from 30 to 250 ms (52, 53). These differences in delay times significantly influence how each type of feedback is utilized in motor control. However, despite the longer delays for visual feedback, it often provides critical information regarding the task goals. Therefore, effective motor control typically requires the integration of both visual and proprioceptive feedback to achieve satisfactory performance.

It is known that humans can integrate visual and haptic information for the purpose of perception (54) and are able to utilize this to guide the generation of movement (55). However, it has also been shown that during movement, healthy humans primarily rely on dynamic estimates of hand motion derived from limb afferent feedback, even when visual information about limb movement is accessible (56). It has been proposed that this is the case because the nervous system considers both sensory variances and temporal delays to achieve optimal multisensory integration and feedback control. Therefore, there is a bias toward using proprioceptive signals rather than the visual signals, due to the former’s shorter latency. Interestingly, our study also shows clear differences in the contextual effect of passive and lead-in movements. Here the effect of visual-only lead-in movements decays faster than that of passive lead-in movements. These differences may also arise due to the longer delays in visual processing, which could mean that we weigh this information less in our estimates of current state.

### The Relevance of Visual and Proprioceptive Feedback

As well as differences in time delays between different modalities of sensory input, other sources of uncertainty might arise due to the phenomena being sensed. Visual feedback depends on observations made in the environment and is affected by external environmental conditions, such as illumination levels, which can change rapidly and unpredictably. Importantly, visual feedback not only corresponds to signals from arm movements, but also from other movements in the environment. Consequently, much visual information may only be indirectly relevant for motor control purposes. As a general principle, it is known that exposure to a more variable range of experiences influences and widens generalization (57). Thus, a broader range of visual experience is consistent with the wider and shallower generalization pattern seen for visual lead-in movements.

In contrast, proprioceptive information arises internally within the body, making it directly related to arm movement. It is also less susceptible to external environmental changes, making it more task-relevant and reliable. Additionally, there is a greater inherent processing delay in the visual pathway, and visual information may arise from multiple observations in the environment. By comparison, the proprioceptive pathway has a shorter inherent delay and relates more directly to the state of the arm. As the time delay increases, uncertainty in the relevance of feedback also increases, but this effect is greater for visual input than for proprioceptive input. This hypothesis helps explain why the directional tuning observed for visual contextual lead-in movements was considerably wider than that of passive movements (16, 18). In the visual condition, there is more uncertainty about the prior movement direction due to the longer processing delay and the observational uncertainty arising from the nature of the environment. We propose that this increased observational uncertainty and longer delays in visual feedback explain the clear difference in the strength of contextual cues with the visual lead-in. The increasing dwell times produce a much weaker integration into the predictive controller than for passive movements.

## Author Contributions

LAH, ISH, and DWF conceived and designed the study. ISH implemented the study on the vBOT, and LAH conducted the data collection. Both LAH and ISH carried out the data analysis on the new visual and passive lead-in data from the current study. ISH carried out the re-analysis of the previous data from a prior study that looked at the effect of dwell time on active lead-in movement (15). LAH and ISH wrote the initial draft of the manuscript. All three authors, LAH, ISH, and DWF, reviewed and edited the manuscript.

## Acknowledgements

Financial support for LAH was provided by the Engineering and Physical Sciences Research Council (EPSRC), and The University of Plymouth. Financial support for ISH was provided by The University of Plymouth. We thank John Welsh for invaluable technical support. During the preparation of this work, the last author (ISH) utilized ChatGPT-4o for proofreading.

## References

1. Shadmehr R, Mussa-Ivaldi FA. Adaptive representation of dynamics during learning of a motor task. J Neurosci 14: 3208–3224, 1994. doi: 10.1523/JNEUROSCI.14-05-03208.1994.

2. Thoroughman KA, Shadmehr R. Learning of action through adaptive combination of motor primitives. Nature 407: 742–747, 2000. doi: 10.1038/35037588.

3. Nozaki D, Kurtzer I, Scott SH. Limited transfer of learning between unimanual and bimanual skills within the same limb. Nat Neurosci 9: 1364–1366, 2006. doi: 10.1038/nn1785.

4. Shadmehr R, Brashers-Krug T. Functional stages in the formation of human long-term motor memory. J Neurosci Off J Soc Neurosci 17: 409–419, 1997. doi: 10.1523/JNEUROSCI.17-01-00409.1997.

5. Krakauer JW, Mazzoni P. Human sensorimotor learning: Adaptation, skill, and beyond. Curr Opin Neurobiol 21: 636–644, 2011. doi: 10.1016/j.conb.2011.06.012.

6. Zarahn E, Weston GD, Liang J, Mazzoni P, Krakauer JW. Explaining Savings for Visuomotor Adaptation: Linear Time-Invariant State-Space Models Are Not Sufficient. J Neurophysiol 100: 2537–2548, 2008. doi: 10.1152/jn.90529.2008.

7. Sing GC, Smith MA. Reduction in Learning Rates Associated with Anterograde Interference Results from Interactions between Different Timescales in Motor Adaptation. PLOS Comput Biol 6: e1000893, 2010. doi: 10.1371/journal.pcbi.1000893.

8. Hamel R, Lepage J, Bernier P. Anterograde interference emerges along a gradient as a function of task similarity: A behavioural study. Eur J Neurosci 55: 49–66, 2022. doi: 10.1111/ejn.15561.

9. Gandolfo F, Mussa-Ivaldi FA, Bizzi E. Motor learning by field approximation. Proc Natl Acad Sci 93: 3843–3846, 1996. doi: 10.1073/pnas.93.9.3843.

10. Krakauer JW, Ghilardi MF, Ghez C. Independent learning of internal models for kinematic and dynamic control of reaching. Nat Neurosci 2: 1026–1031, 1999. doi: 10.1038/14826.

11. Karniel A, Mussa-Ivaldi FA. Does the motor control system use multiple models and context switching to cope with a variable environment? Exp Brain Res 143: 520–524, 2002. doi: 10.1007/s00221-002-1054-4.

12. Caithness G, Osu R, Bays P, Chase H, Klassen J, Kawato M, Wolpert DM, Flanagan JR. Failure to consolidate the consolidation theory of learning for sensorimotor adaptation tasks. J Neurosci Off J Soc Neurosci 24: 8662–8671, 2004. doi: 10.1523/JNEUROSCI.2214-04.2004.

13. Howard IS, Wolpert DM, Franklin DW. The effect of contextual cues on the encoding of motor memories. J Neurophysiol 109: 2632–44, 2013. doi: 10.1152/jn.00773.2012.

14. Brashers-Krug T, Shadmehr R, Bizzi E. Consolidation in human motor memory. Nature 382: 252– 255, 1996. doi: 10.1038/382252a0.

15. Howard IS, Ingram JN, Franklin DW, Wolpert DM. Gone in 0.6 Seconds: The Encoding of Motor Memories Depends on Recent Sensorimotor States. J Neurosci 32: 12756–12768, 2012. doi: 10.1523/JNEUROSCI.5909-11.2012.

16. Howard I, Franklin D. Neural Tuning Functions Underlie Both Generalization and Interference. PloS One 10: e0131268, 2015. doi: 10.1371/journal.pone.0131268.

17. Sarwary M, Stegeman D, Selen L, Medendorp P. Generalization and transfer of contextual cues in motor learning. J Neurophysiol 114: jn.00217.2015, 2015. doi: 10.1152/jn.00217.2015.

18. Howard I, Franklin D. Adaptive tuning functions arise from visual observation of past movement. Sci Rep 6, 2016. doi: 10.1038/srep28416.

19. Howard I, Franklin S, Franklin D. Asymmetry in kinematic generalization between visual and passive lead-in movements are consistent with a forward model in the sensorimotor system. PLOS ONE 15: e0228083, 2020. doi: 10.1371/journal.pone.0228083.

20. Gippert M, Leupold S, Heed T, Howard IS, Villringer A, Nikulin VV, Sehm B. Prior Movement of One Arm Facilitates Motor Adaptation in the Other. J Neurosci 43: 4341–4351, 2023. doi: 10.1523/JNEUROSCI.2166-22.2023.

21. Howard IS, Franklin S, Franklin DW. Kernels of Motor Memory Formation: Temporal Generalization in Bimanual Adaptation. J Neurosci 44, 2024. doi: 10.1523/JNEUROSCI.0359-24.2024.

22. Franklin S, Leib R, Dimitriou M, Franklin D. Congruent visual cues speed dynamic motor adaptation. J Neurophysiol 130, 2023. doi: 10.1152/jn.00060.2023.

23. Howard I, Ford C, Cangelosi A, Franklin D. Active lead-in variability affects motor memory formation and slows motor learning. Sci Rep 7, 2017. doi: 10.1038/s41598-017-05697-z.

24. Howard IS, Ingram JN, Wolpert DM. A modular planar robotic manipulandum with end-point torque control. J Neurosci Meth 181: 199–211, 2009. doi: 10.1016/j.jneumeth.2009.05.005.

25. Oldfield RC. The assessment and analysis of handedness: The Edinburgh inventory. Neuropsychologia 9: 97–113, 1971. doi: 10.1016/0028-3932(71)90067-4.

26. Proud K, Heald JB, Ingram JN, Gallivan JP, Wolpert DM, Flanagan JR. Separate motor memories are formed when controlling different implicitly specified locations on a tool. J Neurophysiol 121: 1342–1351, 2019. doi: 10.1152/jn.00526.2018.

27. Scheidt R, Reinkensmeyer D, Conditt M, Rymer W, Mussa-Ivaldi F. Persistence of Motor Adaptation During Constrained, Multi-Joint, Arm Movements. J Neurophysiol 84: 853–62, 2000. doi: 10.1152/jn.2000.84.2.853.

28. Sadeghi M, Ingram J, Wolpert D. Adaptive coupling influences generalization of sensorimotor learning. PLOS ONE 13: e0207482, 2018. doi: 10.1371/journal.pone.0207482.

29. Orschiedt J, Franklin DW. Learning context shapes bimanual control strategy and generalization of novel dynamics. PLOS Comput Biol 19: e1011189, 2023. doi: 10.1371/journal.pcbi.1011189.

30. Flash T, Hogan N. The coordination of arm movements: an experimentally confirmed mathematical model. J Neurosci 5: 1688–1703, 1985. doi: 10.1523/JNEUROSCI.05-07-01688.1985.

31. Dorf RC, Bishop RH. Modern Control Systems. 7th ed. USA: Addison-Wesley Longman Publishing Co., Inc., 1995.

32. Åström KJ, Murray R. Feedback Systems: An Introduction for Scientists and Engineers, Second Edition. Princeton University Press, 2021.

33. Milner TE, Franklin DW. Impedance control and internal model use during the initial stage of adaptation to novel dynamics in humans. J Physiol 567: 651–664, 2005. doi: 10.1113/jphysiol.2005.090449.

34. Smith MA, Ghazizadeh A, Shadmehr R. Interacting Adaptive Processes with Different Timescales Underlie Short-Term Motor Learning. PLOS Biol 4: e179, 2006. doi: 10.1371/journal.pbio.0040179.

35. Burdet E, Osu R, Franklin DW, Milner TE, Kawato M. The central nervous system stabilizes unstable dynamics by learning optimal impedance. Nature 414: 446–449, 2001. doi: 10.1038/35106566.

36. Franklin DW, Osu R, Burdet E, Kawato M, Milner TE. Adaptation to stable and unstable dynamics achieved by combined impedance control and inverse dynamics model. J Neurophysiol 90: 3270– 3282, 2003. doi: 10.1152/jn.01112.2002.

37. JASP Team, Version 0.19.1. JASP. 2024.

38. Forano M, Schween R, Taylor JA, Hegele M, Franklin DW. Direct and indirect cues can enable dual adaptation, but through different learning processes. J Neurophysiol 126: 1490–1506, 2021. doi: 10.1152/jn.00166.2021.

39. Howard IS, Wolpert DM, Franklin DW. The value of the follow-through derives from motor learning depending on future actions. Curr Biol 25: 397–401, 2015. doi: 10.1016/j.cub.2014.12.037.

40. Sheahan HR, Franklin DW, Wolpert DM. Motor planning, not execution, separates motor memories. Neuron 92: 773–9, 2016. doi: 10.1016/j.neuron.2016.10.017.

41. Goodale MA, Milner AD. Separate visual pathways for perception and action. Trends Neurosci 15: 20–25, 1992. doi: 10.1016/0166-2236(92)90344-8.

42. Grill-Spector K, Malach R. The human visual cortex. Annu Rev Neurosci 27: 649–77, 2004. doi: 10.1146/annurev.neuro.27.070203.144220.

43. DiCarlo JJ, Zoccolan D, Rust NC. How Does the Brain Solve Visual Object Recognition? Neuron 73: 415–434, 2012. doi: 10.1016/j.neuron.2012.01.010.

44. Saunders JA, Knill DC. Humans use continuous visual feedback from the hand to control fast reaching movements. Exp Brain Res 152: 341–352, 2003. doi: 10.1007/s00221-003-1525-2.

45. Smeets J, Brenner E. Fast corrections of movements with a computer mouse. Spat Vis 16: 365–376, 2003. doi: 10.1163/156856803322467581.

46. Franklin DW, Wolpert DM. Computational mechanisms of sensorimotor control. Neuron 72: 425– 442, 2011. doi: 10.1016/j.neuron.2011.10.006.

47. Proske U, Gandevia SC. The Proprioceptive Senses: Their Roles in Signaling Body Shape, Body Position and Movement, and Muscle Force. Physiol Rev 92: 1651–1697, 2012. doi: 10.1152/physrev.00048.2011.

48. Pruszynski JA, Kurtzer I, Scott SH. The long-latency reflex is composed of at least two functionally independent processes. J Neurophysiol 106: 449–459, 2011. doi: 10.1152/jn.01052.2010.

49. Moon KM, Kim J, Seong Y, Suh B-C, Kang K, Choe HK, Kim K. Proprioception, the regulator of motor function. BMB Rep 54: 393–402, 2021. doi: 10.5483/BMBRep.2021.54.8.052.

50. Sainburg RL, Poizner H, Ghez C. Loss of proprioception produces deficits in interjoint coordination. J Neurophysiol 70: 2136–2147, 1993. doi: 10.1152/jn.1993.70.5.2136.

51. Brenner E, van Straaten CAG, de Vries AJ, Baas TRD, Bröring KM, Smeets JBJ. How the timing of visual feedback influences goal-directed arm movements: delays and presentation rates. Exp Brain Res 241: 1447–1457, 2023. doi: 10.1007/s00221-023-06617-6.

52. Miall RC, Jackson J. Adaptation to visual feedback delays in manual tracking: Evidence against the Smith Predictor model of human visually guided action. Exp Brain Res Exp Hirnforsch Expérimentation Cérébrale 172: 77–84, 2006. doi: 10.1007/s00221-005-0306-5.

53. Cameron BD, de la Malla C, López-Moliner J. The role of differential delays in integrating transient visual and proprioceptive information. Front Psychol 5: 50, 2014. doi: 10.3389/fpsyg.2014.00050.

54. Ernst MO, Banks MS. Humans integrate visual and haptic information in a statistically optimal fashion. Nature 415: 429–433, 2002. doi: 10.1038/415429a.

55. Körding KP, Ku S, Wolpert DM. Bayesian integration in force estimation. J Neurophysiol 92: 3161– 3165, 2004. doi: 10.1152/jn.00275.2004.

56. Crevecoeur F, Munoz DP, Scott SH. Dynamic Multisensory Integration: Somatosensory Speed Trumps Visual Accuracy during Feedback Control. J Neurosci Off J Soc Neurosci 36: 8598–8611, 2016. doi: 10.1523/JNEUROSCI.0184-16.2016.

57. Raviv L, Lupyan G, Green SC. How variability shapes learning and generalization. Trends Cogn Sci 26: 462–483, 2022. doi: 10.1016/j.tics.2022.03.007.

